# High-fat diet rewires gut host-microbiota relationship to derive worse outcomes following *Clostridioides difficile* infection

**DOI:** 10.64898/2026.01.05.697779

**Authors:** Archit Kumar, NTQ Nhu Nguyen, Martin O’Brien, Kimberly Vendrov, Alexander Standke, Hannah Ruppel, Ivy Earl, Ingrid L Bergin, Vincent B Young, Raymond Yung

## Abstract

*Clostridioides difficile* is a leading cause of hospital-acquired enteric infections, with outcomes ranging from mild diarrhea to life-threatening pseudomembranous colitis and death. Dietary modulations can influence the course of antibiotic-induced *C. difficile* infection (CDI), and high-fat diets are associated with increased mortality. Here, we profiled host innate immune cells, cecal microbial composition, and the host transcriptome of normal diet (ND) and high-fat diet (HFD)-fed C57BL/6 male and female mice following experimental CDI during the acute and recovery phases. HFD-fed male mice exhibited delayed disease onset but increased mortality following CDI. During acute infection, HFD-fed males showed increased monocyte infiltration and expansion of CD103⁺CD11b⁻ and CD103⁻CD11b⁺ dendritic cell subsets; transcriptomic modules were enriched for antigen presentation and signatures consistent with T cell activation, coincident with enrichment of *Escherichia* and *Lactococcus*. In contrast, during recovery, HFD-fed males and females displayed sustained neutrophil and monocyte infiltration, depletion of eosinophils (prominent in males), tissue-resident Tim4⁺/CD206⁺ macrophage subsets and ILC3s, increased expression of *Tlr4* and endothelial activation markers, downregulation of metabolic pathways, and failure to restore cecal microbial diversity. Collectively, these data indicate that high-fat feeding disrupts resolution programs after CDI and promotes progression to severe disease, with the most pronounced susceptibility in males.

## 1. Introduction

*Clostridioides difficile* (C. difficile), a Gram-positive, spore-forming obligate anaerobe, is a leading cause of nosocomial infections, affecting nearly half a million individuals and contributing to approximately 30,000 deaths annually in the United States^1,2^. Moreover, 20–30% of patients experience recurrent infection due to failure of antibiotic treatment. After colonizing the gut*, C. diffic*ile secretes toxins A (TcdA) and B (TcdB), which cause epithelial damage and trigger an inflammatory response, resulting in clinical manifestations ranging from mild diarrhea to pseudomembranous colitis and death^3–5^.

The gut microbiota confers colonization resistance to *C. difficile* infection (CDI), in part by modulating bile acid composition^6,7^. Microbial derived enzymes convert primary bile acids (cholic acid, β-muricholate, and chenodeoxycholic acid), synthesized by the liver, into secondary bile acids (hyodeoxycholate, ursodeoxycholate, lithocholate, deoxycholate, and ω-muricholate) in the gut, thereby limiting *C. difficile* colonization and growth^7–11^. Commensal microbes play a crucial role in maintaining gut homeostasis; however, antibiotic-induced gut dysbiosis increases the risk of *C. difficile* colonization in the gut^6^. Moreover, restoration of gut homeostasis by fecal microbiota transfer (FMT) has been shown to successfully treat CDI and recurrent CDI^12^.

Dietary changes have been shown to regulate gut microbial diversity and modulate the host response during CDI^13–15^. Diets low in fiber and high in fat and/or protein are associated with worse CDI outcomes, whereas high-fiber, low-protein, high microbiota-accessible carbohydrates (MACs) diets correlate with better survival and reduced disease severity^13,16–22^.

Investigating the impact of dietary changes on the host and gut microbiota is important for developing effective prevention strategies against CDI. Recent studies exploring the effects of a high-fat diet (HFD) on antibiotic-induced CDI in mouse models have demonstrated modulation of host inflammation, bile acids, and gut microbiota, and these changes have been associated with severe inflammation and increased mortality following CDI^19–21^. Although bile acid and microbiota changes in HFD-fed mice during CDI are well characterized, the innate immune landscape and host transcriptional changes during CDI under HFD remain to be fully explored. Here, using an antibiotic-induced CDI model, we profiled myeloid cells and innate lymphoid cells (ILCs) by multi-color spectral flow cytometry and performed weighted gene co-expression network analysis (WGCNA) of the host transcriptome alongside microbiota profiling in normal diet (ND)-fed and HFD-fed male and female mice. We demonstrate that HFD-fed male mice exhibit increased mortality and persistent inflammation, characterized by sustained infiltration of neutrophils and monocytes and loss of eosinophils and tissue-resident Tim4^+^ macrophages during the recovery phase. RNA-seq reveals temporal changes in gene expressions associated with antigen presentation, T cell response, and type I interferon signaling during acute infection. During recovery, upregulation of *Tlr4* and endothelial receptors *Tie1* and *Pecam1*, together with downregulation of oxidative phosphorylation and glutathione metabolism, may sustain inflammation and impede recovery in HFD-fed mice during CDI.

## 2. Results

### 2.1 High-fat diet worsens outcomes following CDI in mice

To assess the effect of a high-fat diet (HFD) on the outcome of *C. difficile* infection (CDI), 4–6-week-old male and female C57BL/6 mice were fed a diet containing 42% kilocalories (kcal) from fat for 12 weeks. Mice maintained on a normal/chow diet (ND) containing 13% kcal from fat for the same duration served as controls. Both male and female HFD-fed mice had significantly higher baseline body weight than ND-fed counterparts; as expected, male mice in general had higher body weight than female mice (Figure S1a). Mice were then given a single intraperitoneal injection of clindamycin (10 mg/kg), were challenged 24 hours later with 10³ spores of *C. difficile* strain VPI 10643 and were monitored for the duration of the study (Figure 1a).

**Figure 1.**
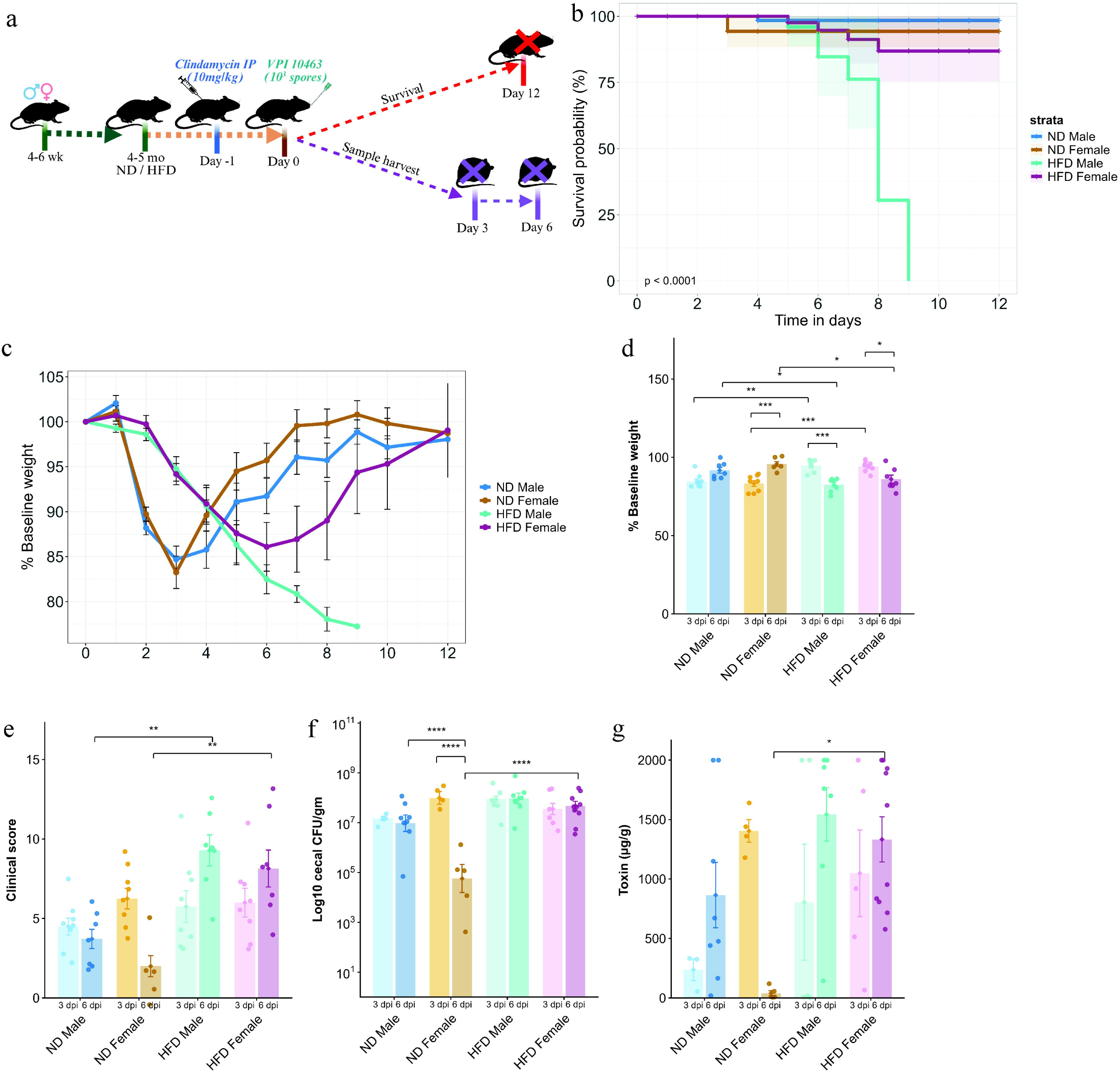
HFD worsens clinical outcomes following CDI. (a) Schematic of the CDI model used for survival and functional studies in male and female mice fed either normal diet (ND) or high-fat diet (HFD) for 12 weeks before infection. (b) Survival probabilities and (c) body weight change over 12 days following CDI. Bar plots showing (d) weight loss and (e) clinical scores (n = 8); (f) Bacterial load and (g) Toxin A concentration in cecal contents obtained from ND- and HFD-fed male and female mice at 3 and 6 dpi following CDI (n = 3-10). Data are presented as mean ± SEM. Statistical significance was assessed by two-way ANOVA followed by Tukey’s HSD post hoc test. p < 0.05 was considered statistically significant. (*) p < 0.05, (**) p < 0.01, (***) p < 0.001, (****) p < 0.0001.

HFD-fed male mice displayed a delayed onset of overt clinical signs and exhibited lower bacterial colonization at 1 day post infection (dpi) compared with ND-fed male mice (Figure S1b,c). ND-fed *C. difficile*–infected male and female mice developed signs of clinical infection within 48 hours post challenge, with maximum weight loss, peak clinical scores, and the first mortality events observed at 3 dpi among ND-fed female mice (Figure 1b-d, S1c). HFD-fed mice showed reduced survival following CDI, and all HFD-fed male mice succumbed to infection by 9 dpi (Figure 1b). ND-fed male and female mice showed maximum weight loss at 3 dpi relative to their baseline, whereas HFD-fed male and female mice exhibited maximum weight loss at 6dpi compared to ND-fed male and female mice respectively (Figure 1c,d). No significant differences in clinical scores, bacterial load, and fecal toxin levels were observed between ND-fed and HFD-fed mice at 3 dpi; however, HFD-fed mice had significantly worse clinical scores, higher bacterial load, and toxin levels at 6 dpi compared to ND-fed counterparts (Figure 1e-g). ND-fed female mice exhibited better signs of recovery at 6dpi in terms of weight gain, clinical scores, bacterial load, and toxin levels (Figure 1b-g and S1c). Interestingly, we observed clear sex-related differences during the recovery phase of CDI, although some mortality occurred among ND-fed female mice during the acute phase, most recovered rapidly following infection, whereas HFD-fed male mice failed to recover and ultimately succumbed to CDI.

### 2.2 HFD-fed mice exhibited severe and prolonged inflammation following CDI

Histopathological analysis was performed to quantify the inflammation and tissue damage in cecal sections from ND- and HFD-fed male and female mice at 3 and 6 dpi following CDI. Notably, histopathology scores mirrored the clinical observations described in the previous section. At 3 dpi, ND-fed male and female mice, as well as HFD-fed female mice, exhibited marked edema, inflammation, and epithelial damage, resulting in high histopathology summary scores (Figure 2a). In contrast, HFD-fed male mice displayed significantly less epithelial damage, lower summary scores, fewer inflammatory cells in the submucosa, and a reduced submucosal area compared to ND-fed male mice at 3dpi (Figure 2a-c). HFD-fed male mice also had significantly less edema and lower summary scores compared to HFD-fed female mice at 3 dpi following CDI (Figure 2a). Representative images are shown in Figure 2d.

**Figure 2.**
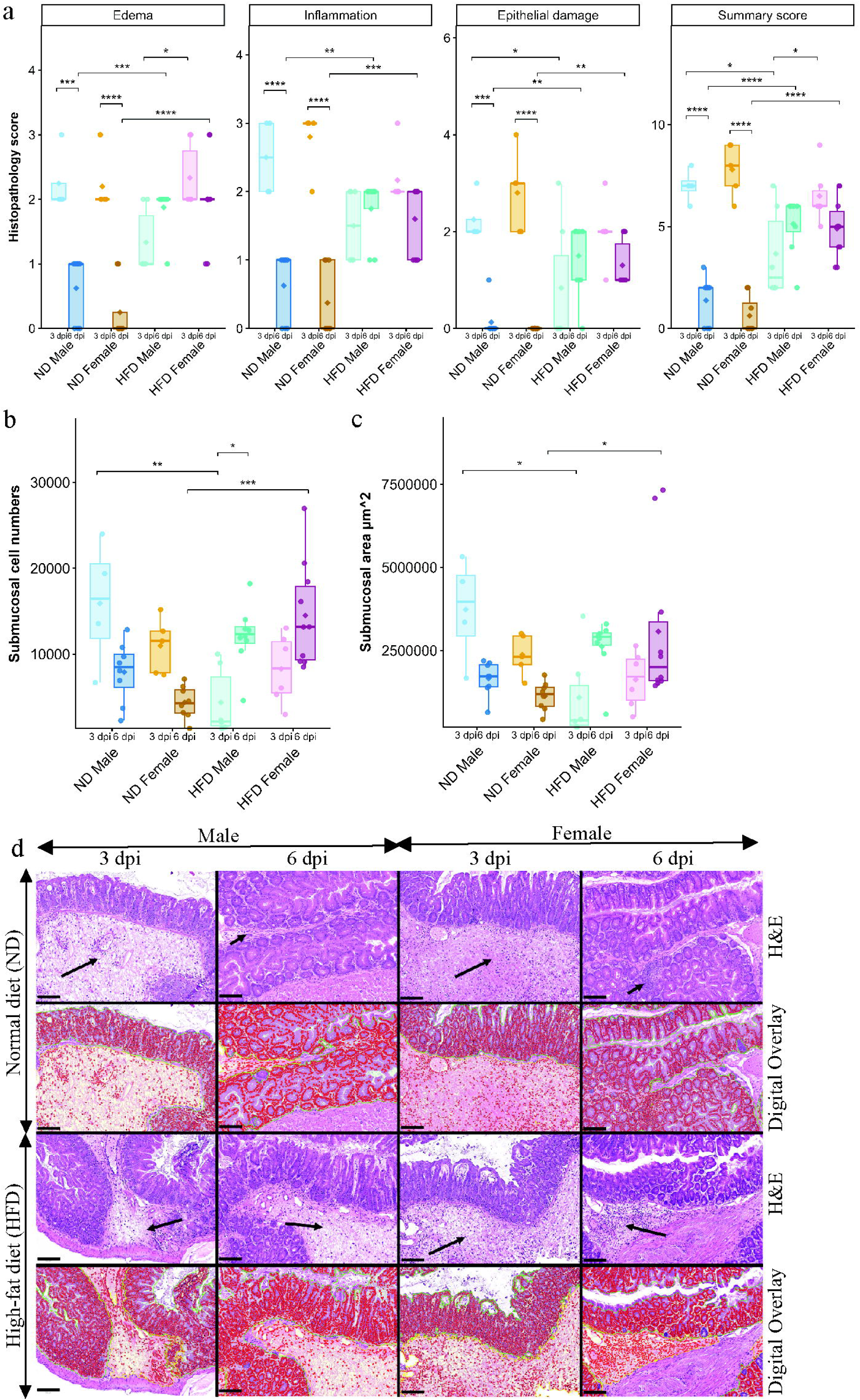
HFD-fed mice had severe cecal inflammation and edema following CDI. (a) Cecal histopathology scores for edema, inflammation, epithelial damage, and total summary scores; (b) submucosal cell counts; (c) submucosal area in ND- and HFD-fed male and female mice at 3 and 6 dpi following CDI (n = 4-10). Histopathology was assessed in a blinded manner. Data are presented as mean ± SEM. Statistical significance was assessed by two-way ANOVA followed by Tukey’s HSD post hoc test; p < 0.05 was considered statistically significant. (d) Representative images of hematoxylin and eosin (H&E)-stained sections and corresponding digital overlays of cecal samples. Long arrows indicate marked submucosal edema and inflammation, and short arrows indicate less severe edema and inflammation. In the digital overlay images, enumerated cells (red) consist of inflammatory cells in the submucosa (yellow outline) and both inflammatory and epithelial cells in the mucosa (green outline). Scale bars: 100 µm. (*) p < 0.05, (**) p < 0.01, (***) p < 0.001, (****) p < 0.0001.

By 6dpi, inflammation and cecal injury had significantly resolved in ND-fed male and female mice compared with 3 dpi (Figure 2a). In HFD-fed male and female mice, edema, inflammation, epithelial damage, and summary scores remained elevated during the recovery phase relative to ND-fed counterparts (Figure 2a). Consistent with this, both the number of inflammatory cells in the submucosa and submucosal area remained higher in HFD-fed mice (Figure 2b,c; representative images in Figure 2d). Together, these findings indicate that HFD feeding is associated with sustained inflammatory cell infiltration and prolonged cecal injury following CDI. Moreover, the lower histopathology summary scores observed in HFD-fed male mice at 3 dpi highlight a distinct transient response in this group, which was no longer evident at 6 dpi.

### 2.3 Altered immune compositions in the colon of HFD-fed mice following CDI

To investigate immunological changes during the acute and recovery phases of CDI, immune cells were isolated from the colonic lamina propria and analyzed by spectral flow cytometry to identify myeloid cells (CD11b⁺), dendritic cells (DCs; MHC-II⁺CD11c⁺), and innate lymphoid cell subsets (ILCs; CD127⁺CD90⁺). The gating strategy used is shown in Figure S2. At baseline, HFD-fed male mice exhibited significant increase in CD206⁻CD11c⁺ macrophages compared with ND-fed male mice and HFD-fed female mice. In contrast, both HFD-fed male and female mice showed a significant reduction in CD206⁺CD11c⁻ and Tim4⁺CD4⁺ macrophages compared to sex-matched ND-fed controls (Figure S3a). No significant baseline differences in DCs were observed between ND-fed and HFD-fed mice (Figure S3b). Collectively, these baseline data indicate that HFD is associated with phenotypic changes among myeloid cells, characterized by an increase in inflammatory macrophages and a decline in wound-healing macrophage subsets.

Acute CDI resulted in a reduction in several innate cell populations, including eosinophils, macrophages, DCs, and ILC3s (Figure S4c,d and Figure S5a). Among myeloid cells, neutrophils were the dominant infiltrating population following CDI, followed by monocytes (Figure S4c). Total CD11b⁺ infiltration did not differ significantly by diet or sex; however, ND-fed female mice exhibited reduced CD11b⁺ infiltration during the recovery phase (Figure S4a). At 3 dpi, no major differences were detected in neutrophils, eosinophils, Tim4⁺CD4⁺ macrophages, CD206⁻CD11c⁺ macrophages, or CD206⁺CD11c⁻ macrophages between HFD-fed and ND-fed mice (Figure 3a; Figure S4b). Notably, HFD-fed male mice exhibited increased monocyte infiltration compared with ND-fed male mice and HFD-fed female mice at 3 dpi. By 6 dpi, both HFD-fed males and females showed significantly increased neutrophil and monocyte infiltration, together with a marked decline in Tim4⁺CD4⁺ macrophages compared with ND-fed males and females. Declines in CD206⁻CD11c⁺ and CD206⁺CD11c⁻ macrophage populations were also observed in HFD-fed mice at 6 dpi compared to ND-fed mice (Figure S4b). Moreover, HFD-fed male mice had significantly lower eosinophil levels than ND-fed male mice at 6 dpi. Among ND-fed female mice, neutrophils significantly declined from 3 dpi to 6 dpi, accompanied by increased eosinophils and macrophage populations (including Tim4⁺ and CD206⁺ subsets) at 6 dpi. Over the same interval, ND-fed male mice and HFD-fed female mice showed a significant increase in CD206⁻CD11c⁺ macrophages (Figure 3a; Figure S4b).

**Figure 3.**
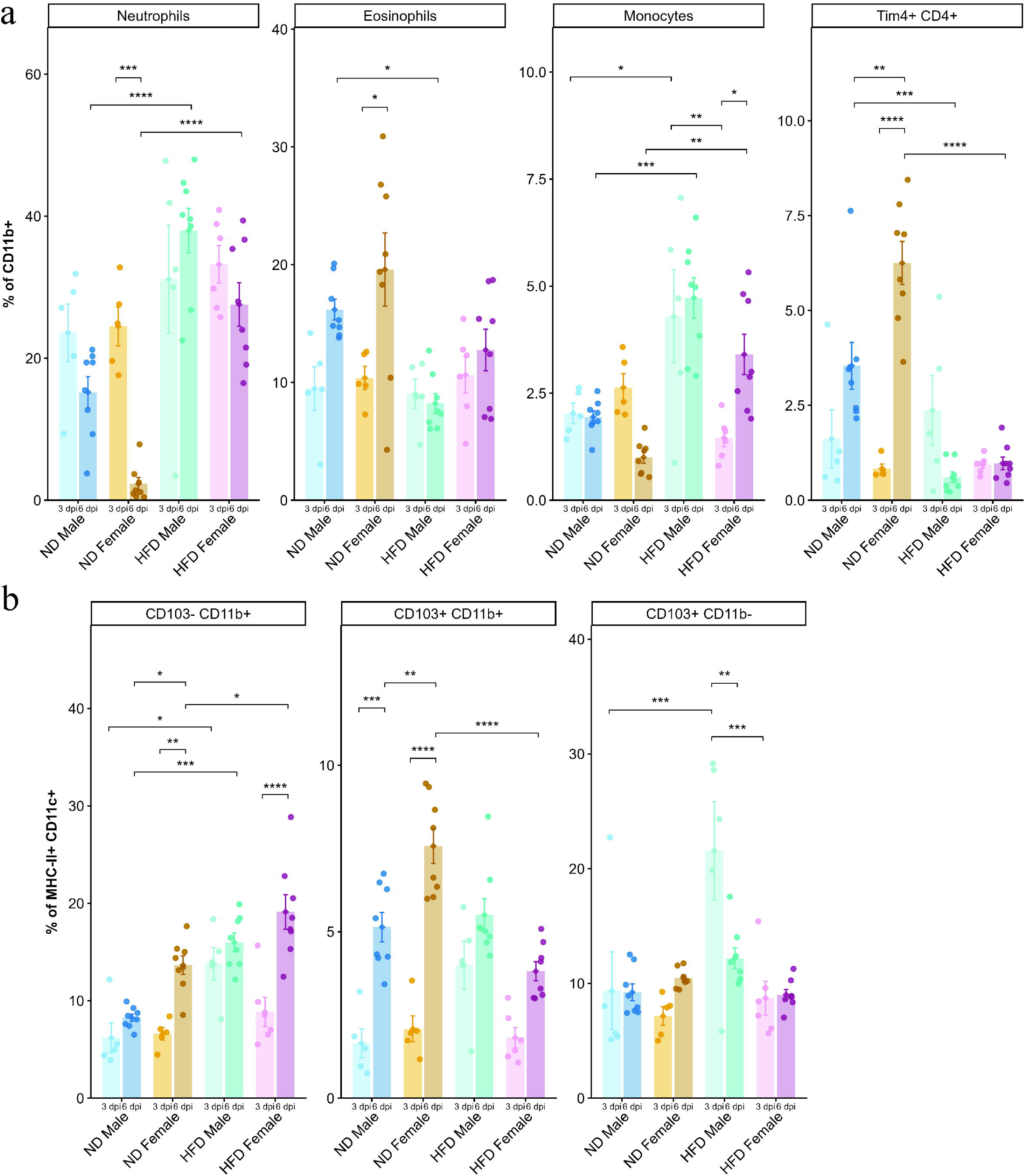
HFD altered colonic innate immune composition following CDI. Bar plots showing the frequencies of (a) CD11b⁺ myeloid cells (neutrophils (Ly6G^+^), eosinophils (Siglec-F^+^, monocytes (Ly6C^+^) and tissue-resident Tim4⁺CD4⁺ macrophages); (b) dendritic cell subsets (MHC-II^+^ CD11c^+^; CD103^-^CD11b^+^, CD103^+^CD11b^+^, and CD103^+^CD11b^-^) in the colonic lamina propria isolated from ND- and HFD-fed male and female mice at 3 and 6 dpi following CDI (n = 5-8). Data are presented as mean ± SEM. Statistical significance was determined by two-way ANOVA followed by Tukey’s HSD post hoc test. p < 0.05 was considered statistically significant. (*) p < 0.05, (**) p < 0.01, (***) p < 0.001, (****) p < 0.0001.

Within the DC compartment, HFD-fed male mice exhibited increased frequencies of CD103⁻CD11b⁺ DCs (type 2 conventional DCs; cDC2) compared with ND-fed male mice at 3 dpi; by 6 dpi, CD103⁻CD11b⁺ cDC2 frequencies were increased in both HFD-fed males and females relative to ND-fed mice (Figure 3b). HFD-fed male mice also had significantly higher frequencies of CD103⁺CD11b⁻ DCs (type 1 conventional DCs; cDC1) than ND-fed male mice and HFD-fed female mice at 3 dpi, followed by a decline at 6 dpi (Figure 3b). CD103⁺CD11b⁺ DCs (cDC2) showed a trend toward increased frequencies from 3 dpi to 6 dpi across all groups; at 6 dpi, ND-fed female mice had higher frequencies than HFD-fed female mice (Figure 3b).

HFD-fed mice exhibited decline in ILC1s with a significant reduction in ILC1s in HFD-fed female mice compared to ND-fed female mice (Figure S3c). No significant baseline differences were observed in ILC2s or ILC3s between ND- and HFD-fed mice (Figure S3c). Following CDI, expansion of ILC1s was primarily observed in ND-fed male and female mice at 3 dpi, whereas expansion of ILC3s was observed at 6 dpi in ND-fed mice (Figure S5a). Consistent with this pattern, ND-fed female mice had significantly higher ILC1 frequencies than HFD-fed female mice at 3 dpi, and ND-fed males and females exhibited significantly higher ILC3 frequencies than HFD-fed counterparts at 6 dpi (Figure S5c). Conversely, both HFD-fed males and females exhibited higher ILC2 frequencies than their ND-fed counterparts at 3 dpi, which subsequently declined by 6 dpi (Figure S5c). Although diet-related differences in ILC cytokine production were limited, ND-fed female mice showed a significant increase in IFNγ-producing ILC1s and IL-13-producing ILC2s at 6 dpi relative to 3 dpi (Figure S5d). HFD-fed mice did not exhibit defects in IL-22–producing ILC3s during the acute phase. At 6 dpi, ND-fed male mice had higher frequencies of IL-22–producing ILC3s than ND-fed female mice and HFD-fed male mice (Figure S5d).

Overall, HFD-fed mice exhibited prolonged infiltration of neutrophils and monocytes together with depletion of eosinophils, and tissue-resident Tim4⁺ and CD206⁺ macrophages, and ILC3s at 6 dpi, a pattern consistent with sustained inflammation and impaired transition to a recovery phase. Moreover, the increase in CD103⁺CD11b⁻ and CD103^-^CD11b^+^ DCs at 3 dpi CDI in HFD-fed male mice suggests enhanced potential for lymphocyte priming and involvement during the acute phase of infection^23^.

### 2.4 HFD-fed mice had reduced diversity and delayed recovery of the gut microbiota following CDI

After demonstrating changes in the innate immune compartment following CDI, we next examined bacterial diversity in cecal samples from ND-fed and HFD-fed mice at baseline, after clindamycin treatment (mock), and following CDI at 3 dpi and 6 dpi. Principal coordinate analysis (PCoA) indicated that the microbial diversity of antibiotic-treated (mock) HFD-fed male and female mice differed from those of ND-fed mice at both 3 dpi and 6 dpi (Figure 4a).

**Figure 4.**
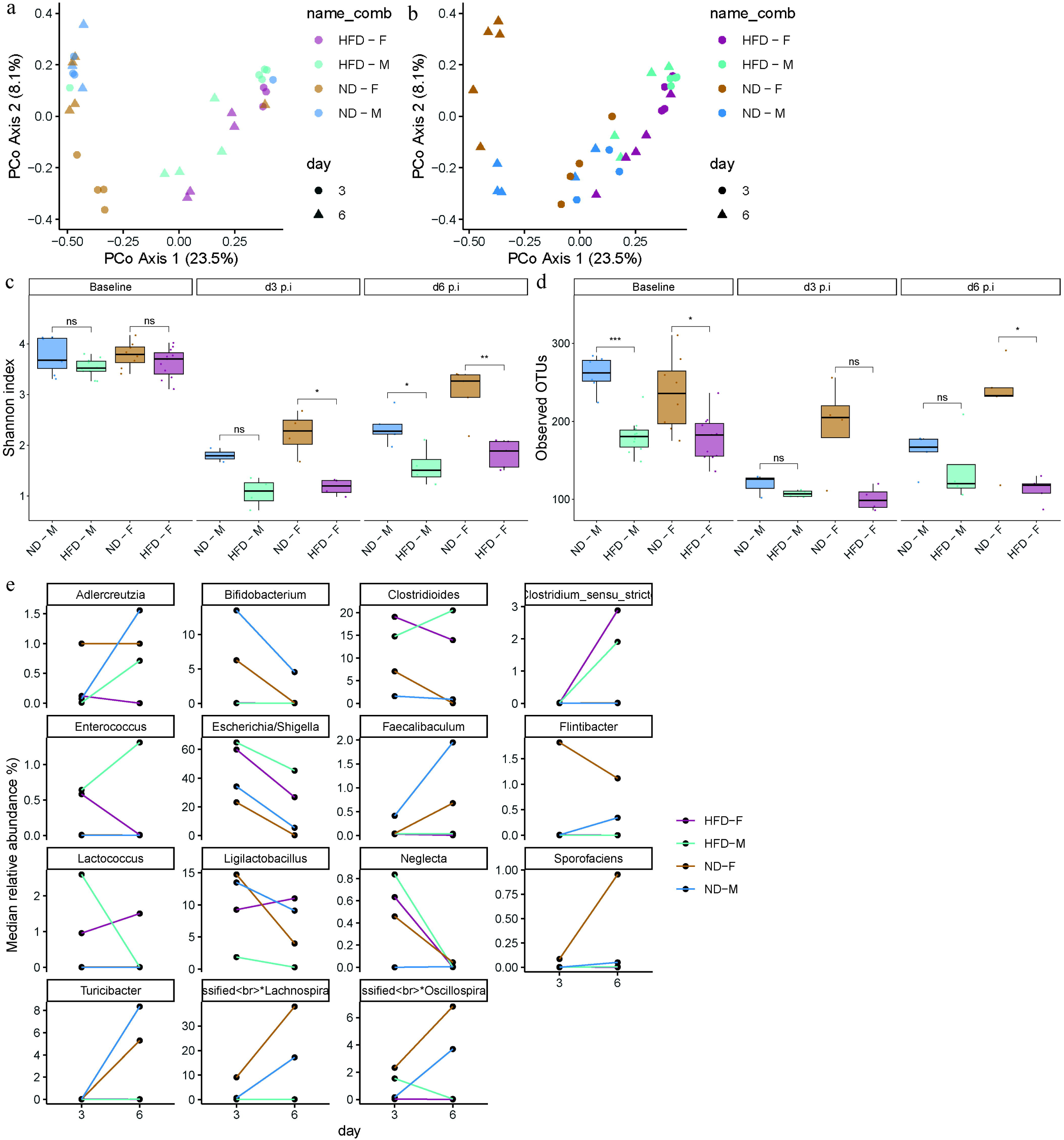
HFD-fed mice exhibited loss of gut microbial diversity before and after CDI. Principal coordinate analysis (PCoA) of fecal microbiota from (a) antibiotic-treated (mock) and (b) CDI ND- and HFD-fed male and female mice at 3 dpi and 6 dpi. Alpha diversity measured using (c) Shannon index and (d) observed operational taxonomic units (OTUs) in ND- and HFD-fed male and female mice at baseline, and CDI (3 and 6 dpi) time points. (e) Line plots showing the relative abundance of significantly altered taxa in ND- and HFD-fed male and female mice at 3 dpi and 6 dpi following CDI. Data is presented as median. Statistical significance was determined by Wilcoxon test. p < 0.05 was considered statistically significant. (*) p < 0.05, (**) p < 0.01, (***) p < 0.001, (****) p < 0.0001.

Although diet- and sex-dependent shifts were observed in both male and female mice at 3 dpi and 6 dpi following CDI, ND-fed female mice exhibited a distinct pattern and showed the greatest divergence at 6 dpi (Figure 4b). HFD-fed male and female mice had a less diverse microbiota at baseline, as reflected by lower Shannon index values and reduced operational taxonomic unit (OTU) richness compared with ND-fed male and female mice (Figure 4c,d). As expected, bacterial diversity declined in all groups after antibiotic treatment and CDI at 3 dpi, with HFD-fed male and female mice showing the most pronounced loss of diversity compared with their ND-fed counterparts during infection period (Figure 4c,d). By 6 dpi, ND-fed female mice exhibited the greatest resilience, with partial recovery of microbial richness, whereas HFD-fed male and female mice still had reduced bacterial richness.

Taxonomic analysis revealed that HFD-fed mice showed increased relative abundance of *Escherichia*, *Clostridioides*, *Enterococcus*, and *Lactococcus*, whereas ND-fed mice had higher abundance of *Akkermansia*, *Ligilactobacillus*, and *Bifidobacterium* at 3 dpi following CDI (Figure 4d, S6). By 6 dpi, along with increased bacterial richness, HFD-fed mice also exhibited higher abundance of *Enterobacter* and *Paenibacillus*, whereas ND-fed mice showed increased abundance of *Turicibacter*, *Lactobacillus*, *Limosilactobacillus*, *Bifidobacterium*, *Flintibacter*, and *Faecalibaculum*, taxa that may contribute to recovery from CDI in ND-fed mice^24–29^.

Overall, these analyses indicate that HFD-fed mice harbor a less diverse gut microbiota that is further depleted by CDI and recovers more slowly. The relative lack of beneficial taxa in HFD-fed mice may reduce colonization resistance and impair recovery, thereby contributing to more severe outcomes following CDI.

### 2.5 HFD-fed mice exhibited distinct transcriptional changes following CDI

To characterize how diet and sex shape host responses to CDI, we profiled cecal gene expression by bulk RNA-seq in ND-fed and HFD-fed male and female mice at 3 dpi and 6 dpi, including mock-infected controls for each condition. We used DESeq2 to model the effects of diet, sex, infection, and time point on gene expression. Principal component analysis (PCA) showed a clear separation of CDI versus mock samples along the first principal component (PC1, 63% of variance), and a further separation of 3 dpi and 6 dpi samples along PC2 (15% of variance) (Figure 5a). To determine how HFD modified transcriptional responses during CDI, we tested a three-way interaction between diet, treatment, and time point, focusing on changes induced during the recovery phase (6 dpi). This analysis identified 3,758 differentially expressed genes (DEGs) with log₂ fold change ≥ 0.5 and FDR < 0.05. Hierarchical clustering of these DEGs revealed that HFD-fed male mice showed distinct temporal expression patterns following CDI compared with ND-fed mice (Figure S7a). Clustering analysis identified three major gene-expression patterns (Figure S7b). One group of genes (cluster 1) was predominantly expressed at early time points and was enriched for cell-cycle–related pathways, consistent with strong proliferative responses during acute infection. A second cluster (cluster 2) was preferentially expressed during the recovery phase and was enriched for fatty acid and amino acid metabolic pathways, suggesting activation of metabolic programs. A third cluster (cluster 3) was expressed mainly in HFD-fed male mice and was enriched for T-cell and B-cell signaling pathways.

**Figure 5.**
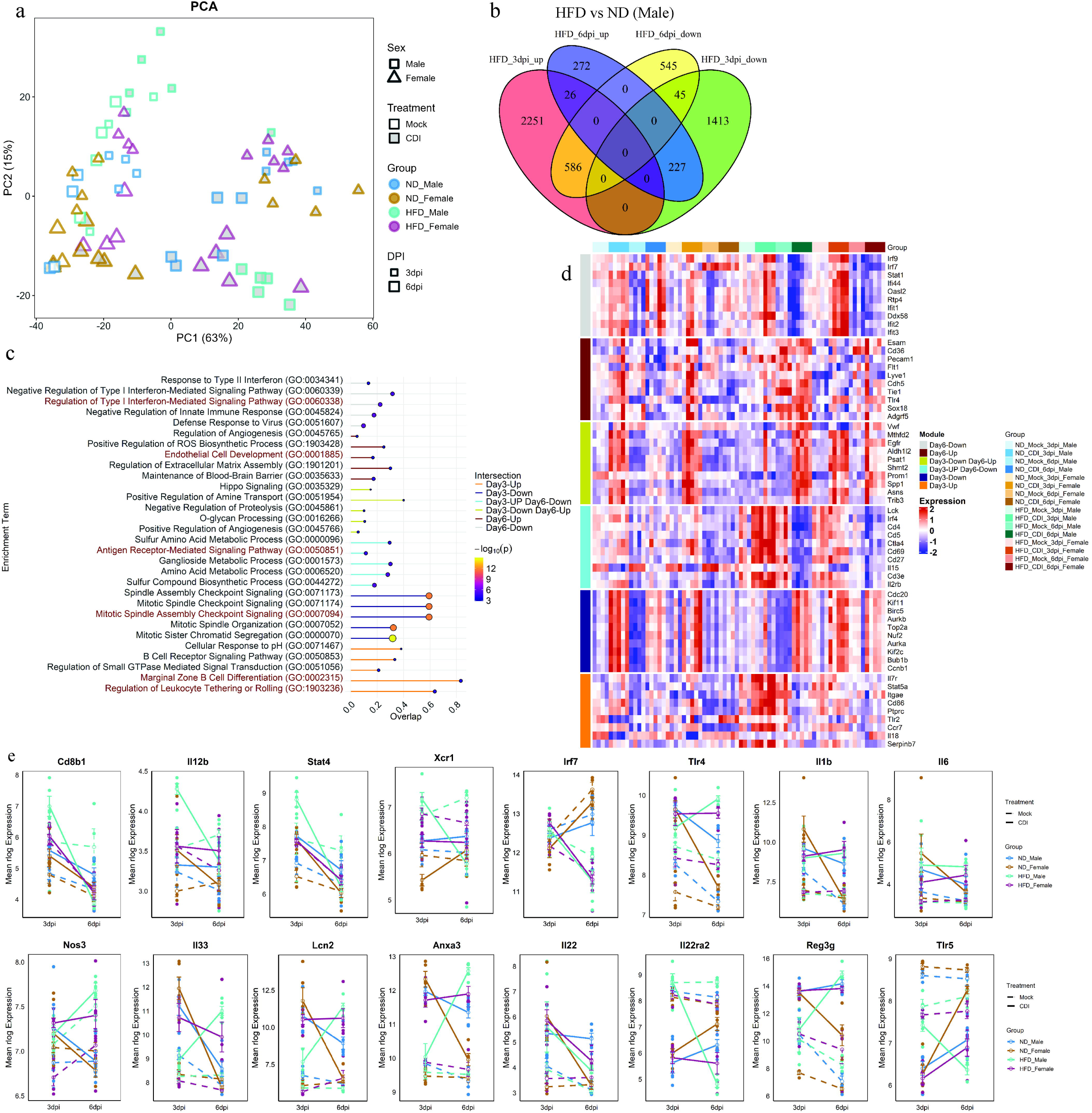
HFD-fed male mice showed temporal changes in gene expression following CDI. (a) Principal component analysis (PCA) of RNA-seq data from cecal tissue obtained from ND- and HFD-fed male and female mice at 3 and 6 dpi following CDI. Antibiotic-treated samples are referred to as mock. Point size indicates dpi, shape indicates sex and grey-filled symbols indicate HFD-fed mice (n = 4-5). (b) Venn diagram of differentially expressed genes (DEGs; log₂ fold change ≥ 0.5 and adjusted p < 0.05) in HFD- versus ND-fed male mice at 3 dpi and 6 dpi. (c) Gene Ontology (GO) analysis showing the top 5 enriched terms for DEGs from each Venn intersection using Enrichr; node size and color represent FDR values. (d) Heatmap showing the expression profiles of the top 5 hub genes from each intersection (identified using CytoHubba) across all groups. (e) Line plots showing log-normalized (rlog) expression levels of selected genes in ND- and HFD-fed male and female mice at 3 dpi and 6 dpi following CDI, including the mock group. Data are presented as mean ± SEM.

Whereas ND-fed mice downregulated cluster 1 and upregulated cluster 2 genes during recovery (6 dpi versus 3 dpi), HFD-fed mice, particularly males, exhibited the opposite pattern along with downregulated cluster 3 at 6 dpi.

As HFD-fed male mice displayed a distinct transcriptional profile and the most pronounced phenotypic changes following CDI, we next performed pairwise comparisons between HFD-fed and ND-fed infected male mice at 3 dpi and 6 dpi using a group-based design. These comparisons yielded 4,548 DEGs at 3 dpi and 1,702 DEGs at 6 dpi. Venn intersection analysis of these DEGs was used to identify genes uniquely regulated in HFD-fed male mice at 3 dpi and 6 dpi (Figure 5b). Expression pattern analysis showed that genes upregulated in HFD-fed male mice at 3 dpi often displayed the opposite behavior at 6 dpi and vice versa (Supplementary Figure S7a-h). Gene Ontology (GO) enrichment analysis of these DEG sets is presented as dot plots, and expression levels of representative hub genes for selected pathways are shown as heat maps (Figure 5c,d).

Functional analysis of 2,251 genes uniquely upregulated at 3 dpi and 586 genes upregulated at 3 dpi and downregulated at 6 dpi in HFD-fed male mice revealed enrichment for leukocyte migration, particularly involving B cells, T cells, and dendritic cells, as well as antigen presentation pathways. These signatures were associated with increased expression of T cell receptor and co-stimulatory/inhibitory genes (e.g., *Cd3e*, *Cd4*, *Cd27*, *Il7r*, *Ctla4*), and signaling molecules (*Lck*, *Stat4*, *Stat5a, Nfkb1*) (Figure 5d, and S9). Consistent with our flow cytometry data, HFD-fed mice showed upregulation of genes characteristic of cDC1, including *Xcr1*, *Id2*, *Itgae* (CD103), *Cd40*, *Cd86* and *Irf8* at 3 dpi (Figure 5d,e and S9). In addition, increased expression of *Cd4*, *Cd8b1*, *Ifng*, *Ccl5*, *Il12b*, *Il16, Stat4*, *Tcf7*, and *Lef1* suggested a cDC1-driven T-cell response (Figure 5d,e, and S9). HFD-fed male mice exhibited increased expression of Nfkb1 and Rela, while showing expression levels of *Il1b*, *Il6*, *Il17a*, *Tnf*, *Cxcl9, Cxcl10*, Ccl2, and Ccl7 comparable to ND-fed male mice at 3 dpi (Figure 5e, and S9). Il22 expression profiles mirrored the IL-22–producing ILC3 data, with no significant differences between groups at 3 dpi, although ND-fed male mice showed a trend toward increased IL-22 production at 6dpi (Figure 5e, 2c). Interesting, HFD-fed male mice had increased transcript levels of decoy receptor *Il22ra2* and decreased levels of anti-microbial peptide *Reg3g* at 3 dpi following CDI (Figure 5e).

Genes uniquely downregulated in HFD-fed male mice at 3 dpi (1,413 DEGs) were enriched for cell-cycle regulation, including *Ccnb1*, *Top2a*, *Kif11*, and *Cdc20* (Figure 5d). A subset of genes (227 DEGs) that were downregulated at 3 dpi and upregulated at 6 dpi was enriched for amine transport and angiogenesis pathways (*Vwf*). At 6 dpi, functional analysis of genes upregulated in HFD-fed male mice indicated involvement of endothelial cells and angiogenesis (*Cdh5*, *Pecam1*, *Tie1, Esam, Sox18, Nos3*) and regulation by *Tlr4*, whereas genes uniquely downregulated at 6 dpi (545 DEGs) were associated with interferon signaling, including *Stat1*, *Irf7*, *Irf9*, and *Oasl2* (Figure 5d,e).

Notably, both HFD-fed male and HFD-fed female mice exhibited increased expression of genes involved in inflammation, such as *Il33*, *Ccl6*, *Cd5l*, *Tnf*, *Ly6g*, *Ly6c1*, *Cxcl1*, *Cxcl2*, *Lcn2*, and *Anxa3* at 6 dpi, whereas ND-fed female mice showed downregulation of these inflammatory genes and ND-fed male mice had intermediate expression levels (Figure 5e, and S9). ND-fed mice also gained expression of *Tlr5* at 6 dpi, which has been proposed to confer protection against CDI^30^, whereas HFD-fed male mice showed downregulation of *Tlr5* at the same time point (Figure S9). Overall, these analyses indicate that HFD-fed male mice exhibit a unique transcriptional response to CDI, characterized by enhanced DCs, and T cell-related responses at 3 dpi and sustained upregulation of inflammatory and endothelial-associated genes at 6 dpi, in contrast to the more resolving metabolism-associated programs observed in ND-fed mice.

### 2.6 WGCNA linked impaired metabolic process and altered microbiota to poor recovery in HFD-fed mice

Given the distinct transcriptional changes observed in HFD-fed mice during CDI, we next asked which genes correlated most strongly with diet, sex, infection, and time. We applied weighted gene co-expression network analysis (WGCNA) to group genes into modules with similar expression profiles and related their eigengenes to experimental traits. WGCNA resolved 13 color-coded modules that were significantly correlated (positively or negatively; FDR < 0.05) with one or more groups (Figure 6a). Modules MEblue, MEred, and MEgreenyellow were positively associated with the acute phase (3 dpi) in ND-fed mice. In HFD-fed males, expression in MEblue shifted temporally, while MEgreenyellow remained significantly correlated at 3 dpi. Modules that were enriched during the recovery phase (6 dpi) were largely confined to ND-fed females (MEmagenta, MEroyalblue) and showed no positive correlation with HFD groups at 6 dpi. Although MEturquoise was only weakly correlated with recovery (FDR < 0.1), it was significantly negatively correlated with the acute phase in ND-fed mice, suggesting changes that may facilitate recovery after CDI. MElightcyan correlated positively only in males (ND- and HFD-fed). HFD-fed males showed distinct positive correlations with MEbrown, MElightgreen, and MEcyan at 3dpi, while HFD-fed females showed a positive correlation with MEdarkgreen at 3 dpi, which was weakly positive (FDR < 0.1) in HFD-fed males.

**Figure 6.**
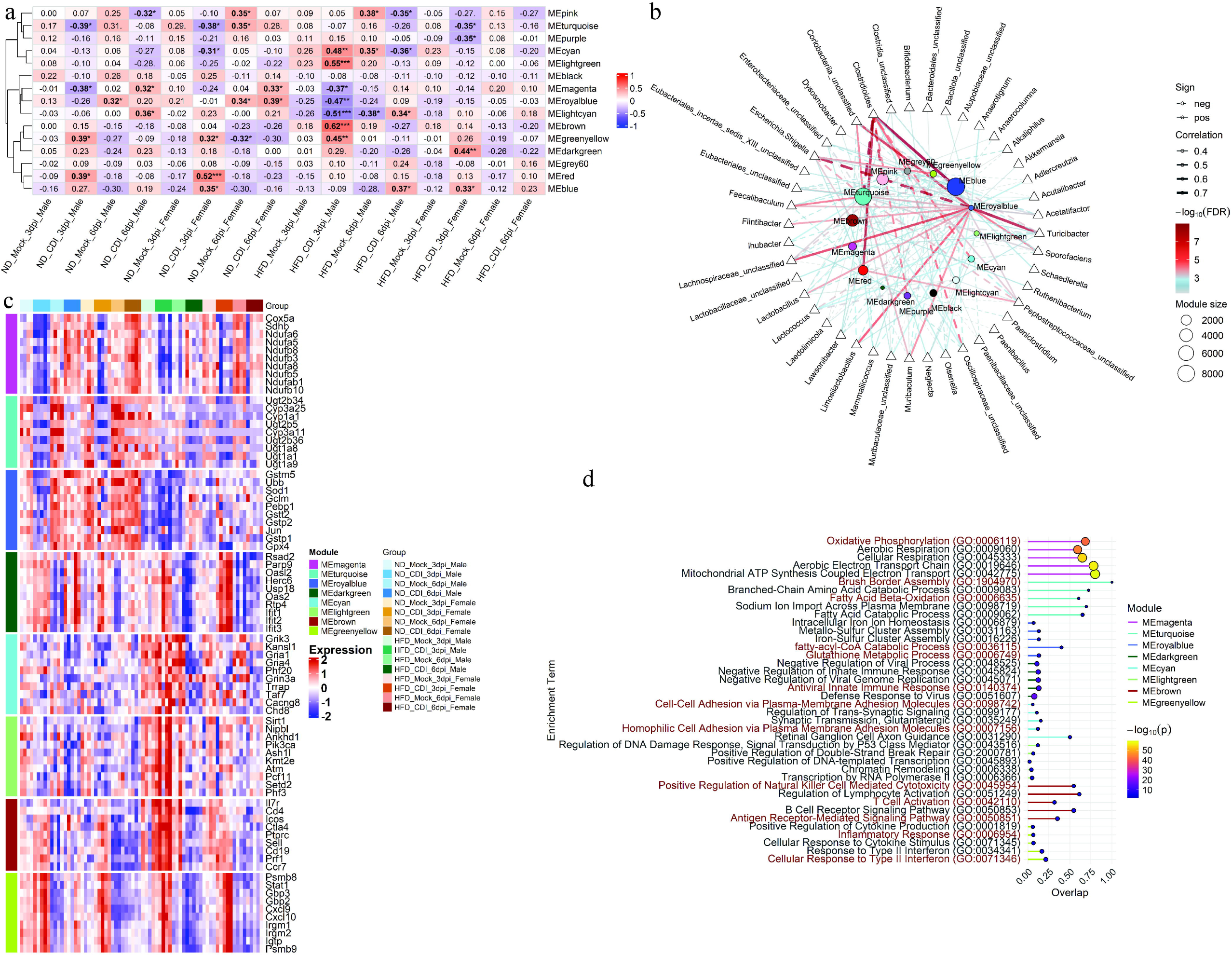
HFD-fed mice exhibited defects in metabolic processes and loss of microbial diversity during the recovery phase following CDI. Weighted gene co-expression network analysis (WGCNA) was performed on RNA-seq data from 16 groups comprising ND- and HFD-fed male and female mice at 3 dpi and 6 dpi following CDI, along with mock groups. (a) Heatmap showing correlations between module eigengenes and traits. (b) Network showing correlations between module eigengenes and bacterial taxa. Line color represents FDR, line thickness represents correlation strength and line style indicates positive (solid) or negative (dotted) correlations; node size represents the number of genes in each module. (c) Heatmap showing expression profiles of the top 5 hub genes from each module (identified using CytoHubba). (d) GO analysis showing the top 5 enriched terms for genes in each module using Enrichr; node size and color represent FDR values. For WGCNA, P values were adjusted for multiple testing using the Benjamini–Hochberg false discovery rate (FDR), and FDR < 0.05 was considered significant. (.) FDR < 0.1; (*) FDR < 0.05; (**) FDR < 0.01; (***) FDR < 0.001; (****) FDR < 0.0001.

Considering microbial taxa can modulate host transcription and influence CDI outcomes^28,29,31–35^, we next tested whether module eigengenes correlated with cecal bacterial taxa. Using Spearman correlations between eigengenes and taxa abundances, MEblue, MEred, and MEgreenyellow were positively associated with *Clostridioides* and also correlated with Escherichia and *Peptostreptococcaceae* (Figure 6b, S10). The male-associated MElightcyan module correlated positively with *Clostridioides* and *Turicibacter*. Recovery-phase modules MEroyalblue and MEturquoise, prominent in ND-fed females at 6 dpi, showed strong positive correlation with taxa *Coriobacteriia*, *Eubacteriales*, *Faecalibaculum*, *Flintibacter, Lachnospiraceae*, *Lactobacillus*, *Limosilactobacillus*, *Muribaculum*, *Sporofaciens*, and *Turicibacter*, and correlated negatively with *Clostridioides* and *Escherichia*. Modules associated with HFD-fed males (MEbrown, MEcyan, MElightgreen) all showed positive correlations with *Lactococcus*; while MEbrown also correlated with *Escherichia* and *Ruthenibacterium*, whereas MEcyan also correlated positively with *Neglecta* and negatively with *Clostridioides*. No significant microbiota correlations were detected for module MEdarkgreen.

We next performed functional enrichment and hub gene identification within modules strongly associated with HFD or with recovery in ND-fed mice (6 dpi) (Figure 6c,d). MEgreenyellow was enriched for interferon-driven inflammatory response (*Stat1*, *Cxcl9*, *Cxcl10*) among HFD-fed and ND-fed mice. MEbrown was enriched for T-cell related (*Il7r*, *Cd4*, *Icos*) and antigen-presentation signaling; its positive association with *Escherichia* and *Lactococcus* is consistent with reports that these taxa can potentiate mucosal immune activation^36,37^. MElightgreen and MEcyan were enriched for chromatin regulation and cell adhesion pathways among HFD males. MEdarkgreen genes were enriched for type I interferon signaling (*Ifit* genes, *Oasl2*, *Rsad2*) and were upregulated in HFD-fed mice of both sexes relative to ND controls during the acute phase following CDI (3 dpi). Recovery-associated modules MEroyalblue, MEturquoise, and MEmagenta were enriched for glutathione metabolism (*Gst* family genes, *Sod1*), fatty acid metabolic pathways, oxidative phosphorylation (*Cox5a*, *Nduf* family genes), and xenobiotic detoxification, including genes such as *Ugt1a1* and *Cyp1a1*.

Overall, HFD-fed mice, particularly males, showed strong positive correlations with modules linked to *Escherichia* and *Lactococcus* and inflammatory/T-cell programs, and type I interferon–associated inflammatory state during acute CDI. In contrast, the recovery phase in ND-fed mice, especially females, was characterized by modules associated with *Faecalibaculum*, *Flintibacter*, *Lactobacillus*/*Limosilactobacillus*, *Muribaculum*, and related taxa, together with enrichment for glutathione metabolism, oxidative phosphorylation, and detoxification programs, features that may promote epithelial repair and resolution. These module–taxa associations are correlative and do not establish causality, but they highlight candidate microbe–host programs that differ between HFD- and ND-fed mice during CDI.

## 3. Discussion

Disruption of commensal gut microbiota is a major risk factor for *C. difficile* infection. Dietary changes can modulate microbial composition and, in turn, affect gut homeostasis. While high-fiber diets have been associated with beneficial outcomes following CDI, diets rich in fat are linked to worse outcomes^13,16,17,19–21^. Identifying how diet shapes host responses during the acute and recovery phases following CDI is therefore important for preventing severe disease. In this study, we found that HFD-fed male mice exhibited a delayed onset of clinical disease but had increased mortality following CDI. The acute phase of CDI in HFD-fed male mice was characterized by increased frequencies of monocytes and CD103^+^ CD11b^-^ DCs (cDC1) and enrichment of cDC1-associated genes, which could lead to T-cell-mediated response. In contrast, during the recovery phase, HFD-fed male mice showed a reduction in eosinophils, Tim4^+^ and CD206^+^ macrophages, downregulation of *Tlr5*, inability to restore commensal microbiota along with downregulation of metabolic pathways, persistent infiltration on neutrophils and monocytes, upregulation of *Tlr4* and angiogenesis- related pathways (*Pecam1*, *Tie1, Esam, Sox18, Vwf*, *Cdh5*), and downregulation of type-I interferon response features which could lead to sustained inflammation and impaired resolution.

HFD-fed mice, notable in males, showed activation of cDC1-mediated T cell response during the acute phase (3 dpi) of the disease following CDI. HFD-fed male mice had a significant increase in CD103+ CD11b- DCs in the colonic lamia propria and showed increased transcript levels of *Itgae* (CD103), *Xcr1*, and *Irf8*, along with *Cd40* and *Cd86* in cecal tissue, suggesting the activation of cDC1 during the acute phase of the CDI. Along with the activation of cDC1 they also showed upregulated expression levels of genes involved in T cell-mediated response, including *Cd4*, *CD8b1*, *Cd3e*, *Cd69*, *Il7r*, *Lck*, *Cd27*, and *Il12b*. HFD-fed female also exhibited increased transcript levels of these genes; however, their expression levels were lower than those of HFD-fed males. Antibiotic-treated HFD-fed male mice (mock) showed similar changes in gene expression; however, given that dietary antigens have a limited role in regulating colonic T cell responses, the HFD-associated enrichment of *Escherichia* and *Lactococcus* is likely to have driven these transcriptional changes^36,37,48–50^ Furthermore, WGCNA identified a gene module associated with antigen-mediated T cell responses (MEbrown) that correlated strongly with HFD-fed mice at 3 dpi following CDI, as well as with *Escherichia* and *Lactococcus*, suggesting that the expansion of these taxa during CDI might have further increased the transcript levels of these genes. While CD103⁺CD11b⁻ DCs (cDC1) have been shown to restrict DSS-induced colitis by inducing T cell–derived IFN-γ^23^, how cDC1-driven T cell responses contribute to CDI pathology in the context of HFD is unclear. In line with the previous report and our transcriptomic data indicate that HFD can promote a Th1-skewed response in the gut^38^ . A shift from Th1 toward Th2/Th17 has been shown to be associated with more severe disease^39^, and Th1 response has been suggested to protect the gut during CDI^40^. We therefore hypothesize that HFD-induced Th1 activation may have non-specifically benefited HFD-fed mice, particularly males, leading to a transient delay in disease onset, reduced colonization at day 1 post infection and less epithelial damage at 3 dpi. Furthermore, increased transcript levels of the decoy receptor *Il22ra2* (IL-22BP), which has been reported to be produced mainly by cDCs and CD4⁺ T cells^41,42^, could have attenuated IL-22 signaling and resulted in reduced *Reg3g* expression^43^ in HFD-fed male mice during the acute phase of infection, thereby predisposing them to more severe tissue damage.

Although the overall proportion of CD11b⁺ myeloid cells during CDI in HFD-fed mice remained unchanged during the recovery phase, marked shifts within myeloid subsets were observed.

Consistent with earlier reports, HFD-fed mice showed increased frequencies of neutrophils and monocytes compared with ND-fed mice^21^. Persistent infiltration of neutrophils and monocytes during the recovery phase (6 dpi) may contribute to the gut damage and mortality observed in HFD-fed mice following CDI^44–46^. Moreover, HFD-fed mice, pronounced in males, exhibited marked upregulation of *Tlr4*, *Tie1*, *Cdh5*, *Sox18*, *Vwf*, *Pecam1*, and *Nos3* during the recovery phase following CDI. TLR4 expressed on myeloid cells recognizes surface layer proteins of *C. difficile* and initiates a downstream inflammatory response^47^. Activation of TLR4 signaling during the acute phase in ND-fed mice could drive an initial inflammatory response that helps bacterial clearance^47,48^, whereas its persistent upregulation in HFD-fed mice could increase the risk of inflammatory gut damage. Moreover, HFD itself increases the expression of *Tlr4* via long-chain saturated fatty acids^49^, as can be seen in HFD-mock mice compared to ND-fed mock mice, suggesting an inflammatory state among HFD-fed mice independent of CDI. Further, HFD-fed mice showed upregulation of endothelial and angiogenesis-related genes *Pecam1*, *Tie1*, *Cdh5*, *Sox18,* and *Nos3,* during the recovery phase which could further aggravate gut inflammation by recruiting additional immune cells^50–53^. While an enrichment of *Firmicutes* has been linked to increased angiogenesis during HFD exposure^54^, and HFD-fed male mice also showed increase in *Enterococcus* at 6 dpi, which could contribute to angiogenesis, we were unable to detect a significant correlation between the two using WGCNA analysis.

In parallel, HFD-fed male mice showed a decline in eosinophils and macrophages during the recovery phase compared with ND-fed mice. While eosinophils have been shown to play a protective role in CDI pathogenesis, CD206⁺ macrophages have also been suggested to contribute to recovery^55,56^. The epithelial alarmins IL-25 and IL-33 have been found to promote eosinophil recruitment during CDI^56,57^ and have also been proposed to modulate CD206⁺ macrophages^55^. We observed upregulation of *Il33* expression during the acute phase (3 dpi) of CDI in ND-fed male and female mice and HFD-fed female mice, whereas HFD-fed male mice showed increased *Il33* expression primarily during the recovery phase (6 dpi). In contrast, ND-fed female mice showed better recovery and displayed increased frequencies of eosinophils, and CD206⁺ and Tim4⁺ macrophages, and IL-13-producing ILC2s during the recovery phase compared with the acute phase. It is therefore plausible that early *Il33* induction during acute CDI in ND-fed female mice might have promoted downstream activation of ILC2s and, eosinophil and macrophage responses during recovery phase^55,57^. Although ND-fed male mice showed a similar trend in eosinophil and macrophage proportions to ND-fed female mice during the recovery phase, the response in females might be skewed by survival bias, as only surviving female mice were analyzed, whereas males still exhibited signs of infection at 6 dpi. By contrast, HFD-fed male mice had a marked decline in eosinophils along with embryonically seeded Tim4⁺ macrophages, implicated in wound healing and gut homeostasis^58,59^, and had a baseline increase in inflammatory CD206^-^ CD11c⁺ macrophages, suggesting that HFD-fed male mice enter infection with a pre-existing pro-inflammatory state in the gut.

Both ND-fed and HFD-fed mice showed upregulation of genes involved in type-I interferon signaling, *Ddx58*, *Ifr7*, *Irf9*, *Stat1*, and *Ifi44,* during the acute phase of CDI. However, HFD-fed male and female mice showed marked decline in the transcript levels of interferon-stimulated genes during the recovery phase following CDI. Type 1 interferon plays an important role in mediating anti-inflammatory effects and in regulating gut homeostasis^60,61^. While expression of type I interferon could be beneficial by limiting inflammation or inducing tolerance^60–62^, their downregulation could aggravate inflammatory response in the gut^63^. HFD-fed mice exhibited worse clinical signs at 6 dpi, the loss of type I interferons could have permitted unrestricted inflammation, as reflected by the persistent high expression of transcripts such as *Il1b*, *Il6*, and *Il17a*. WGCNA indicated that modules enriched for metabolic pathways, including oxidative phosphorylation, glutathione metabolism, and xenobiotic metabolism, showed no positive correlation with HFD-fed mice. By contrast, ND-fed mice, notable in females, displayed strong positive correlations and higher expression of these metabolic modules, which were in turn strongly associated with bacterial taxa such as *Faecalibaculum*, *Flintibacter*, *Lachnospiraceae*, *Lactobacillus*, *Limosilactobacillus*, *Muribaculum*, and *Turicibacter*, taxa that have been implicated in maintaining gut homeostasis and supporting recovery following CDI^29,31,33,35,64–68^. Restoration of metabolic processes in ND-fed mice could have aided in maintaining gut homeostasis and promoting recovery following CDI^69–72^. Among HFD-fed male and female mice, failure to restore microbial diversity and maintain homeostasis during the recovery phase after CDI may be linked to persistent tissue damage and increased mortality.

In summary, our integrative analysis of immune, host transcriptomic, and microbiota profiles reveals that HFD reshapes CDI in a time-dependent manner. The acute phase of CDI in HFD-fed mice is characterized by cDC1-driven T cell response, likely driven in part by *Escherichia* and/or *Lactococcus*, which may contribute to the delayed onset of clinical symptoms and might have affected IL-22 signaling. By contrast, the recovery phase is marked by persistent infiltration of neutrophils and monocytes, increased TLR4 and angiogenesis, failure to engage pro-resolving programs due to loss of eosinophils and CD206⁺/Tim4⁺ macrophages, downregulation of type I interferon and metabolic pathways, and limited restoration of gut microbiota following CDI. Together, these findings provide insights into how HFD could modulate the host response over the course of CDI to exacerbate tissue pathology, impair resolution, and increase mortality.

## 4. Methods

### 4.1 Mice and Diet

Young 4-6 weeks old male and female C57BL/6J mice housed at the University of Michigan were used in this study. Mice were either fed with high-fat diet (HFD) (42% Kcal from fat, Inotiv; TD.88137) or normal diet (ND) (13% Kcal from fat, LabDiet; 5L0D) for 12 weeks. No microbiota normalization was performed between ND-fed and HFD-fed mice. All the experiments were approved by the Unit of Laboratory Animal Medicine, University of Michigan, under animal protocols PRO00010459 and were performed accordingly.

### 4.2 *C. difficile* infection model

Sixteen-eighteen weeks old mice were intraperitoneally (i.p.) injected with clindamycin (10mg/kg) (Sigma). After 24 hours, mice were inoculated with 10^3^ VPI 10463 via oral gavage, while mock mice were given distilled water at the same time. Animals were observed daily for clinical signs such as weight loss, diarrhea, activity, posture, coat and eyes, and nose^73^. The clinical scores were calculated using 4-point scale for diarrhea, activity, posture, coat condition, and appearance of eyes and nose, and mice were euthanized using CO2 inhalation if clinical scores exceeded14 points.

### 4.3 Histopathology

Following euthanasia, cecum samples were harvested from mice at 3 and 6 dpi and fixed in Carnoy’s solution. Samples were processed to paraffin by routine methods, stained using hematoxylin and eosin and subjected to blinded assessment by a board-certified veterinary pathologist at the Unit for Laboratory Animal Medicine Pathology Core. Histological severity scoring performed for edema, inflammation, and epithelial damage, were calculated using a 4-point scale for each parameter. Summary scores represent sum of individual scores for edema, inflammation, and epithelial damage. Quantitative analysis was performed on digitized slide files within the open-source analysis program QuPath v0.6.0^74^. Mucosa and submucosa were manually annotated, and total cell counts were conducted within these regions using a cell count plug-in within software. Lumen and areas of artifact were manually excluded from the region annotations, and the muscular wall was excluded from subsequent analysis

### 4.4 Bacterial load and toxin quantification

To check for *C. difficile* colonization, cecal content collected from euthanized mice was firstly diluted with sterile PBS (Invitrogen). Diluted cecal content was plated on Chrom agar by Spiral plater. The plates were incubated in anaerobic chamber at 37C overnight. Colony forming unit was counted and calculated using ProtoCOL3.

For toxin quantification, cecal content was diluted to 1:1000 in sterile PBS. The procedure has been described previously^75^. Toxin is detected using RTCA system. Briefly, filtered cecal content was incubated with HT-29 cells monolayer, in electrode-lined 96-well plates (E-Plate View 96; ACEA Biosciences) and changes in cell-induced electrical impedance were monitored for 24 hours. The activity of toxins in cecal samples is compared with a standard curve using *C. difficile* toxin A (List Biological Labs).

### 4.5 Microbiota

To investigate the microbiota of ND- and HFD-fed male and female mice at baseline, we collected fecal pellets one day before antibiotic treatment. Microbiota samples after infection were collected from the cecum at 3 and 6 dpi from both antibiotic-treated and *C. difficile* infected mice. Samples were submitted to University of Michigan Microbiome Core for DNA extraction and 16S sequencing based on the protocol by Pat Schloss as described earlier^75,76^.

### 4.6 Isolation of immune cells

Immune cells were isolated from colonic lamina propria as described previously with some modifications^77^. Briefly, colon samples were cut longitudinally into cold DPBS and cleaned. To remove the epithelial cells, colon samples were incubated in a buffer containing HBSS, 0.5mM EDTA, 1mM DTT and 5% FBS in a shaking water bath at 37°C for 15 mins. Samples were vigorously vortexed at an interval of 5 mins for 30 seconds. Tissue samples were rinsed twice with DPBS, minced, and incubated in digestion buffer containing RPMI 1640, 62.5µg/ml Liberase TM (Sigma), 50µg/ml DNase1 (Sigma) and 5%FBS for 45 minutes in a shaking water bath at 37°C. Cells were vigorously vortexed at an interval of 15 mins for 30 seconds. Finally, cells were centrifuged at 600Xg for 6 minutes at 4°C, washed, strained using 70 µm filter and collected in RPMI 1640 with 10% FBS, 2mM L-glutamine, and 1X penicillin/streptomycin.

### 4.7 Flow cytometry

Freshly isolated cells were stained for myeloid cells and innate lymphoid cells (ILCs). To capture intracellular cytokines from ILCs, 1-2 million isolated cells were incubated in culture media containing 1X Brefeldin-A (BioLegend) for 4 hours at 37 °C. For surface staining, cells were first washed using DPBS and stained for live/dead cells using Zombie Aqua (BioLegend). Cells were further washed and incubated with anti-mouse CD16/CD32 TruStain FcX (BioLegend) to block Fc receptors as per manufacturer’s instruction. For myeloid cells staining, cells were incubated with True-Stain Monocyte Blocker (BioLegend) along with Fc block according to manufacturer’s instructions. Following blocking, cells were washed and stained for surface antibodies for 30 minutes. For myeloid cells staining, cells were fixed using BD cytofix fixation buffer (BD Biosciences). For ILCs, cells were fixed, permeabilized using eBioscience™ Foxp3 staining buffer (Thermo) and further stained for cytokines. Cells were acquired in ID7000 (Sony) at University of Michigan Flow Cytometry Core Facility, and unmixed files were analyzed using FlowJo v10.10.0 (FlowJoLLC).

Antibodies used for myeloid cells staining:

CD45 (30-F11, BD Biosciences), CD4 (RM4-5, Thermo), CD19 (6D5, BioLegend), CD3 (17A2, BioLegend), NK1.1 (PK136, BioLegend), CD11b (M1/70, BioLegend), Siglec-F (S17007L, BioLegend), Ly6G (1A8, BioLegend), MHC-II (M5/114.15.2, BD Biosciences), Tim-4 (RMT4-54, BioLegend), CD206 (C068C2, BioLegend), CD64 (X54-5/7.1, BioLegend), Ly6C (HK1.4, BioLegend), CD11c (N418, BD Biosciences), CD103 (2E7, BioLegend).

Antibodies used for ILCs staining:

CD45 (30-F11, BioLegend), CD3 (17A2, BioLegend), CD19 (6D5, BioLegend), Gr-1 (RB6-8C5, BioLegend), F4/80 (BM8, BioLegend), TCRβ (H57-597, BioLegend), TCRγδ (GL3, BioLegend), CD5 (53-7.3, BioLegend), CD8a (53-6.7, BioLegend), CD90.2 (53-2.1, BioLegend), CD127 (A7R34, BioLegend), GATA3 (L50-823, BD Biosciences), T-bet (O4-46, BD Biosciences), RORgt (Q31-378, BD Biosciences), Eomes (Dan11mag, Thermo), IL22 (IL22JOP, Thermo), IL13 (eBio13A, Thermo), IFNγ (XMG1.2, BD Biosciences).

### 4.8 RNAseq

Cecum tissue samples from mock and CDI mice were used for RNAseq analysis. Briefly, tissue samples were processed for RNA isolation using Direct-zol™ RNA Miniprep (Zymo Research). RNA samples were then submitted to University of Michigan Advanced Genomics Core for quality check, library preparation, sequencing and generating count files. Briefly, RNA libraries were prepared using poly-A enrichment and subjected to 151bp paired-end sequencing according to the manufacturer’s protocol (Illumina NovaSeqXPlus, System Suite Version: 1.3.0.39308).

Differential expression analysis was performed using DEseq2 v 1.42.1^78^ using design ∼ Diet*Treatment*DPI*Sex , or ∼Group. Pairwise comparisons computed using ∼Group design were subjected to Log2 fold change shrinkage using apeglm^79^. Genes with Log2 fold change cutoff of ±0.5 and padj < 0,05 were considered as differential expressed genes (DEGs).

Enrichment analysis on DEGs was performed using online tool Enrichr^80^. Hub genes analysis was performed using Maximal Clique Centrality (MCC) method on cytoHubba available on Cytosccape v 3.10.3^81,82^.

### 4.9 Weighted gene co-expression network analysis (WGCNA)

Normalized counts matrix obtained following variance stabilizing transformation were used for the WGCNA analysis. A signed co-expression network was constructed using WGCNA v 1.73 package^83^. Briefly, threshold of R^2^ ≥ 0.89 was used for approximate scale-free topology and softPower = 9 was used for adjacency matrix and TOMsimilarity. Gene modules were identified using dynamic tree cutting with deepSplit = 2 and minimum module size = 50. Further, modules with high similarities were merged based on eigengene correlation of 0.8 using cutHeight = 0.2. Pearson correlation between module eigengene and trait was plotted as heatmap using pheatmap v 1.0.13^84^ and ComplexHeatmap v 2.18.0^85^. Spearman correlation was performed between module eigengene and microbial taxa. All p-values were adjusted for multiple testing using Benjamini–Hochberg false discovery rate (FDR) method.

### 4.10 Statistical analysis

Data is presented as mean ± standard error of mean (SEM). Graphs are generated using ggplot2. Statistical significance for baseline samples was calculated using one-way ANOVA followed by Tukey’s HSD post hoc test using ggpubr^82^. Survival data was analyzed using survminer and ggsurvfit packages^86,87^. Two-way ANOVA followed by Tukey’s HSD post hoc test was used to assess statistical significance among the *C. difficile* infected groups using ggpubr. Adjusted p-value of less than 0.05 was considered significant.

## Authors contribution

Study design: A.K., N.N., V.Y., and R.Y. Experiments: A.K., M.O’B., N.N., K.V. A.S., H.R., and I.E. Data acquisition and analysis: A.K., N.N. and I.L.B. Writing: A.K. Reviewing and editing: A.K., N.N., I.L.B, V.Y. and R.Y. Funding: V.Y. and R.Y.

## Funding

This work was supported by the National Institute of Health, grant number R01AI162787 from the National Institute of Allergy and Infectious Diseases (NIAID).

## Acknowledgements

We thank the Unit for Laboratory Animal Medicine Pathology Core (RRID:SCR_018823, University of Michigan) for performing histopathology; the Microbiome Core (University of Michigan) for DNA extraction and 16S sequencing; and the Advanced Genomics Core (RRID:SCR_026820, University of Michigan) for library preparation and next-generation sequencing. BioRender was used to create schematic figures, and ChatGPT was used for English correction and text clarity.

## Data availability

The RNA-seq data generated in this study have been deposited in the NCBI Gene Expression Omnibus (GEO) under accession GSE313697.

## Notes

### Competing Interest Statement

The authors have declared no competing interest.

## References

1. Lessa, F. C., Winston, L. G., McDonald, L. C., & Emerging Infections Program C. difficile Surveillance Team. Burden of Clostridium difficile infection in the United States. N Engl J Med 372, 2369–2370 (2015).

2. Guh, A. Y. et al. Trends in U.S. Burden of Clostridioides difficile Infection and Outcomes. N Engl J Med 382, 1320–1330 (2020).

3. Carter, G. P. et al. Defining the Roles of TcdA and TcdB in Localized Gastrointestinal Disease, Systemic Organ Damage, and the Host Response during Clostridium difficile Infections. mBio 6, e00551 (2015).

4. Ng, J. et al. Clostridium difficile toxin-induced inflammation and intestinal injury are mediated by the inflammasome. Gastroenterology 139, 542–552, 552.e1–3 (2010).

5. Voth, D. E. & Ballard, J. D. Clostridium difficile toxins: mechanism of action and role in disease. Clin Microbiol Rev 18, 247–263 (2005).

6. Theriot, C. M., Bowman, A. A. & Young, V. B. Antibiotic-Induced Alterations of the Gut Microbiota Alter Secondary Bile Acid Production and Allow for Clostridium difficile Spore Germination and Outgrowth in the Large Intestine. mSphere 1, e00045–15 (2016).

7. Foley, M. H. et al. Bile salt hydrolases shape the bile acid landscape and restrict Clostridioides difficile growth in the murine gut. Nat Microbiol 8, 611–628 (2023).

8. Smits, W. K., Lyras, D., Lacy, D. B., Wilcox, M. H. & Kuijper, E. J. Clostridium difficile infection. Nat Rev Dis Primers 2, 16020 (2016).

9. Theriot, C. M. & Young, V. B. Interactions Between the Gastrointestinal Microbiome and Clostridium difficile. Annu Rev Microbiol 69, 445–461 (2015).

10. Paredes-Sabja, D., Shen, A. & Sorg, J. A. Clostridium difficile spore biology: sporulation, germination, and spore structural proteins. Trends Microbiol 22, 406–416 (2014).

11. Chiang, J. Y. L. Bile acids: regulation of synthesis. J Lipid Res 50, 1955–1966 (2009).

12. Bhat, A. et al. Safety and efficacy of fecal microbiota transplantation versus antibiotics for treating clostridioides difficile infection: systematic review and meta-analysis. Eur J Clin Microbiol Infect Dis (2025) doi:10.1007/s10096-025-05278-3.

13. Castro, M., Silver, H. J., Hazleton, K., Lozupone, C. & Nicholson, M. R. The Impact of Diet on Clostridioides difficile Infection: A Review. J Infect Dis 231, e1010–e1018 (2025).

14. Zmora, N., Suez, J. & Elinav, E. You are what you eat: diet, health and the gut microbiota. Nat Rev Gastroenterol Hepatol 16, 35–56 (2019).

15. Martinez, K. B., Leone, V. & Chang, E. B. Western diets, gut dysbiosis, and metabolic diseases: Are they linked? Gut Microbes 8, 130–142 (2017).

16. Hewlett, K. K. et al. Dietary Fiber Modulates the Window of Susceptibility to Clostridioides difficile Infection. Gastroenterology 169, 970–982 (2025).

17. Fachi, J. L. et al. Fiber- and acetate-mediated modulation of MHC-II expression on intestinal epithelium protects from Clostridioides difficile infection. Cell Host Microbe 33, 235–251.e7 (2025).

18. Bhute, S. S. et al. A High-Carbohydrate Diet Prolongs Dysbiosis and Clostridioides difficile Carriage and Increases Delayed Mortality in a Hamster Model of Infection. Microbiol Spectr 10, e0180421 (2022).

19. Mefferd, C. C. et al. A High-Fat/High-Protein, Atkins-Type Diet Exacerbates Clostridioides (Clostridium) difficile Infection in Mice, whereas a High-Carbohydrate Diet Protects. mSystems 5, e00765–19 (2020).

20. Hazleton, K. Z. et al. Dietary fat promotes antibiotic-induced Clostridioides difficile mortality in mice. NPJ Biofilms Microbiomes 8, 15 (2022).

21. Jose, S. et al. Obeticholic acid ameliorates severity of Clostridioides difficile infection in high fat diet-induced obese mice. Mucosal Immunol 14, 500–510 (2021).

22. Moore, J. H. et al. Defined Nutrient Diets Alter Susceptibility to Clostridium difficile Associated Disease in a Murine Model. PLoS One 10, e0131829 (2015).

23. Muzaki, A. R. B. M. et al. Intestinal CD103(+)CD11b(-) dendritic cells restrain colitis via IFN-γ-induced anti-inflammatory response in epithelial cells. Mucosal Immunol 9, 336–351 (2016).

24. Ni, Y. et al. Bifidobacterium and Lactobacillus improve inflammatory bowel disease in zebrafish of different ages by regulating the intestinal mucosal barrier and microbiota. Life Sci 324, 121699 (2023).

25. Ojima, M. N. et al. Bifidobacterium bifidum Suppresses Gut Inflammation Caused by Repeated Antibiotic Disturbance Without Recovering Gut Microbiome Diversity in Mice. Front Microbiol 11, 1349 (2020).

26. Li, Y. et al. Bifidobacterium breve synergizes with Akkermansia muciniphila and Bacteroides ovatus to antagonize Clostridioides difficile. ISME J 19, wraf086 (2025).

27. Wu, Y. et al. Galactooligosaccharides and Limosilactobacillus reuteri synergistically alleviate gut inflammation and barrier dysfunction by enriching Bacteroides acidifaciens for pentadecanoic acid biosynthesis. Nat Commun 15, 9291 (2024).

28. Li, J. et al. Limosilactobacillus mucosae-derived extracellular vesicles modulates macrophage phenotype and orchestrates gut homeostasis in a diarrheal piglet model. NPJ Biofilms Microbiomes 9, 33 (2023).

29. Cao, Y. G. et al. Faecalibaculum rodentium remodels retinoic acid signaling to govern eosinophil-dependent intestinal epithelial homeostasis. Cell Host Microbe 30, 1295–1310.e8 (2022).

30. Jarchum, I., Liu, M., Lipuma, L. & Pamer, E. G. Toll-like receptor 5 stimulation protects mice from acute Clostridium difficile colitis. Infect Immun 79, 1498–1503 (2011).

31. Foley, M. H. et al. Differential modulation of post-antibiotic colonization resistance to Clostridioides difficile by two probiotic Lactobacillus strains. mBio 16, e0146825 (2025).

32. AbdelKhalek, A. & Narayanan, S. K. Comparison between Symptomatic and Asymptomatic Mice after Clostridioides difficile Infection Reveals Novel Inflammatory Pathways and Contributing Microbiota. Microorganisms 10, 2380 (2022).

33. Rastogi, S. & Singh, A. Gut microbiome and human health: Exploring how the probiotic genus Lactobacillus modulate immune responses. Front Pharmacol 13, 1042189 (2022).

34. Qi, H. et al. Induction of Inflammatory Macrophages in the Gut and Extra-Gut Tissues by Colitis-Mediated Escherichia coli. iScience 21, 474–489 (2019).

35. Seekatz, A. M. et al. Restoration of short chain fatty acid and bile acid metabolism following fecal microbiota transplantation in patients with recurrent Clostridium difficile infection. Anaerobe 53, 64–73 (2018).

36. Nagayama, M. et al. TH1 cell-inducing Escherichia coli strain identified from the small intestinal mucosa of patients with Crohn’s disease. Gut Microbes 12, 1788898 (2020).

37. Kimoto, H., Mizumachi, K., Okamoto, T. & Kurisaki, J.-I. New Lactococcus strain with immunomodulatory activity: enhancement of Th1-type immune response. Microbiol Immunol 48, 75–82 (2004).

38. Luck, H. et al. Regulation of obesity-related insulin resistance with gut anti-inflammatory agents. Cell Metab 21, 527–542 (2015).

39. Yu, H. et al. Cytokines Are Markers of the Clostridium difficile-Induced Inflammatory Response and Predict Disease Severity. Clin Vaccine Immunol 24, e00037–17 (2017).

40. Yang, H. et al. A commensal protozoan attenuates Clostridioides difficile pathogenesis in mice via arginine-ornithine metabolism and host intestinal immune response. Nat Commun 15, 2842 (2024).

41. Pelczar, P. et al. A pathogenic role for T cell-derived IL-22BP in inflammatory bowel disease. Science 354, 358–362 (2016).

42. Martin, J. C. J. et al. Interleukin-22 binding protein (IL-22BP) is constitutively expressed by a subset of conventional dendritic cells and is strongly induced by retinoic acid. Mucosal Immunol 7, 101–113 (2014).

43. Sadighi Akha, A. A., et al. Interleukin-22 and CD160 play additive roles in the host mucosal response to Clostridium difficile infection in mice. Immunology 144, 587–597 (2015).

44. Liao, X. et al. Origin and Function of Monocytes in Inflammatory Bowel Disease. J Inflamm Res 17, 2897–2914 (2024).

45. Huber, A. et al. Olfactomedin-4 + neutrophils exacerbate intestinal epithelial damage and worsen host survival after Clostridioides difficile infection. *bioRxiv* 2023.08.21.553751 (2023) doi:10.1101/2023.08.21.553751.

46. Jose, S. & Madan, R. Neutrophil-mediated inflammation in the pathogenesis of Clostridium difficile infections. Anaerobe 41, 85–90 (2016).

47. Ryan, A. et al. A role for TLR4 in Clostridium difficile infection and the recognition of surface layer proteins. PLoS Pathog 7, e1002076 (2011).

48. Lai, Y.-H. et al. The Role of Toll-Like Receptor-2 in Clostridioides difficile Infection: Evidence From a Mouse Model and Clinical Patients. Front Immunol 12, 691039 (2021).

49. Hu, X. et al. Toll-like receptor 4 is a master regulator for colorectal cancer growth under high-fat diet by programming cancer metabolism. Cell Death Dis 12, 791 (2021).

50. Randi, A. M. & Laffan, M. A. Von Willebrand factor and angiogenesis: basic and applied issues. J Thromb Haemost 15, 13–20 (2017).

51. Savant, S. et al. The Orphan Receptor Tie1 Controls Angiogenesis and Vascular Remodeling by Differentially Regulating Tie2 in Tip and Stalk Cells. Cell Rep 12, 1761–1773 (2015).

52. Rijcken, E. et al. PECAM-1 (CD 31) mediates transendothelial leukocyte migration in experimental colitis. Am J Physiol Gastrointest Liver Physiol 293, G446–452 (2007).

53. Ala, A., Dhillon, A. P. & Hodgson, H. J. Role of cell adhesion molecules in leukocyte recruitment in the liver and gut. Int J Exp Pathol 84, 1–16 (2003).

54. Andriessen, E. M. et al. Gut microbiota influences pathological angiogenesis in obesity-driven choroidal neovascularization. EMBO Mol Med 8, 1366–1379 (2016).

55. Donlan, A. N. et al. IL-13 protects from Clostridioides difficile colitis. Anaerobe 88, 102860 (2024).

56. Buonomo, E. L. et al. Microbiota-Regulated IL-25 Increases Eosinophil Number to Provide Protection during Clostridium difficile Infection. Cell Rep 16, 432–443 (2016).

57. Frisbee, A. L. et al. IL-33 drives group 2 innate lymphoid cell-mediated protection during Clostridium difficile infection. Nat Commun 10, 2712 (2019).

58. De Schepper, S. et al. Self-Maintaining Gut Macrophages Are Essential for Intestinal Homeostasis. Cell 175, 400–415.e13 (2018).

59. Hoeffel, G. et al. Sensory neuron-derived TAFA4 promotes macrophage tissue repair functions. Nature 594, 94–99 (2021).

60. Vasquez Ayala, A., et al. Commensal bacteria promote type I interferon signaling to maintain immune tolerance in mice. J Exp Med 221, e20230063 (2024).

61. Kotredes, K. P., Thomas, B. & Gamero, A. M. The Protective Role of Type I Interferons in the Gastrointestinal Tract. Front Immunol 8, 410 (2017).

62. Snyder, D. T. et al. Type I IFN dependent and independent mechanisms of protection in *C. difficile* infection. The Journal of Immunology 196, 65.27–65.27 (2016).

63. Katakura, K. et al. Toll-like receptor 9-induced type I IFN protects mice from experimental colitis. J Clin Invest 115, 695–702 (2005).

64. Niu, H.-Y. et al. Flavonifractor porci sp. nov. and Flintibacter porci sp. nov., two novel butyrate-producing bacteria of the family Oscillospiraceae. Int J Syst Evol Microbiol 75, 006767 (2025).

65. Song, Z. et al. Sea buckthorn berries alleviate ulcerative colitis via regulating gut Faecalibaculum rodentium-mediated butyrate biosynthesis. Phytomedicine 139, 156490 (2025).

66. Lynch, J. B. et al. Gut microbiota Turicibacter strains differentially modify bile acids and host lipids. Nat Commun 14, 3669 (2023).

67. Pereira, F. C. et al. Rational design of a microbial consortium of mucosal sugar utilizers reduces Clostridiodes difficile colonization. Nat Commun 11, 5104 (2020).

68. Zhang, J. et al. Beneficial effect of butyrate-producing Lachnospiraceae on stress-induced visceral hypersensitivity in rats. J Gastroenterol Hepatol 34, 1368–1376 (2019).

69. Ghosh, S., Moorthy, B., Haribabu, B. & Jala, V. R. Cytochrome P450 1A1 is essential for the microbial metabolite, Urolithin A-mediated protection against colitis. Front Immunol 13, 1004603 (2022).

70. Tubau-Juni, N. et al. Modulation of colonic immunometabolic responses during Clostridioides difficile infection ameliorates disease severity and inflammation. Sci Rep 13, 14708 (2023).

71. Yang, N., Sun, R., Liao, X., Aa, J. & Wang, G. UDP-glucuronosyltransferases (UGTs) and their related metabolic cross-talk with internal homeostasis: A systematic review of UGT isoforms for precision medicine. Pharmacol Res 121, 169–183 (2017).

72. Hayes, J. D., Flanagan, J. U. & Jowsey, I. R. Glutathione transferases. Annu Rev Pharmacol Toxicol 45, 51–88 (2005).

73. Warren, C. A. et al. Amixicile, a novel inhibitor of pyruvate: ferredoxin oxidoreductase, shows efficacy against Clostridium difficile in a mouse infection model. Antimicrob Agents Chemother 56, 4103–4111 (2012).

74. Bankhead, P. et al. QuPath: Open source software for digital pathology image analysis. Sci Rep 7, 16878 (2017).

75. Abernathy-Close, L. et al. Intestinal Inflammation and Altered Gut Microbiota Associated with Inflammatory Bowel Disease Render Mice Susceptible to Clostridioides difficile Colonization and Infection. mBio 12, e0273320 (2021).

76. Kozich, J. J., Westcott, S. L., Baxter, N. T., Highlander, S. K. & Schloss, P. D. Development of a dual-index sequencing strategy and curation pipeline for analyzing amplicon sequence data on the MiSeq Illumina sequencing platform. Appl Environ Microbiol 79, 5112–5120 (2013).

77. Abernathy-Close, L. et al. Aging Dampens the Intestinal Innate Immune Response during Severe Clostridioides difficile Infection and Is Associated with Altered Cytokine Levels and Granulocyte Mobilization. Infect Immun 88, e00960–19 (2020).

78. Love, M. I., Huber, W. & Anders, S. Moderated estimation of fold change and dispersion for RNA-seq data with DESeq2. Genome Biol 15, 550 (2014).

79. Zhu, A., Ibrahim, J. G. & Love, M. I. Heavy-tailed prior distributions for sequence count data: removing the noise and preserving large differences. Bioinformatics 35, 2084–2092 (2019).

80. Kuleshov, M. V. et al. Enrichr: a comprehensive gene set enrichment analysis web server 2016 update. Nucleic Acids Res 44, W90–97 (2016).

81. Chin, C.-H. et al. cytoHubba: identifying hub objects and sub-networks from complex interactome. BMC Syst Biol 8 **Suppl 4**, S11 (2014).

82. Shannon, P. et al. Cytoscape: a software environment for integrated models of biomolecular interaction networks. Genome Res 13, 2498–2504 (2003).

83. Langfelder, P. & Horvath, S. WGCNA: an R package for weighted correlation network analysis. BMC Bioinformatics 9, 559 (2008).

84. Kolde, R. pheatmap: Pretty Heatmaps. R package version 1.0.13. https://github.com/raivokolde/pheatmap.

85. Gu, Z., Eils, R. & Schlesner, M. Complex heatmaps reveal patterns and correlations in multidimensional genomic data. Bioinformatics 32, 2847–2849 (2016).

86. Kassambara, A., Kosinski, M. & Biecek, P. survminer: Drawing Survival Curves using ‘ggplot2’. R package version 0.5.1. https://rpkgs.datanovia.com/survminer/index.html.

87. Sjoberg, D., Baillie, M., Fruechtenicht, M., Haesendonckx, S. & Treis, T. ggsurvfit: Flexible Time-to-Event Figures. R package version 1.2.0. https://github.com/pharmaverse/ggsurvfit.

