## Supplemental Figures for "High-fat diet rewires gut host-microbiota relationship to derive worse outcomes following *Clostridioides difficile* infection"

**Supplementary Figures:**

Figure S1. HFD-fed mice exhibited worse clinical outcomes following CDI. (a) Baseline body weight of male and female mice after 12-weeks of high-fat diet (HFD) feeding, along with normal diet (ND)-fed control (n = 20-24). (b) Bacterial colonization quantified in fecal samples collected after 24 hours *C. difficile* infection (CDI) in ND-fed and HFD-fed male and female mice (n = 8). (c) Clinical scores over the study period in ND-fed and HFD-fed male and female mice following CDI (n = 8). Data are presented as mean ± SEM. Statistical significance was assessed by two-way ANOVA followed by Tukey's HSD post hoc test. $p < 0.05$ was considered statistically significant. (*) $p < 0.05$, (**) $p < 0.01$, (***) $p < 0.001$, (****) $p < 0.0001$.

Figure S2. Flow-cytometry gating strategy used to define (a) myeloid cell populations, (b) innate lymphoid cells (ILCs), and (c) cytokines produced by ILC1s (T-bet$^+$), ILC2s (GATA-3$^+$), and ILC3s (ROR-$\gamma$T$^+$).

Figure S3. HFD feeding affected colonic macrophage populations. Baseline changes in innate immune cell composition in the colonic lamina propria isolated from ND- and HFD-fed male and female mice: (a) myeloid cells, (b) dendritic cells, and (c) innate lymphoid cells (n = 3-4). Data are presented as mean ± SEM. Statistical significance was determined by two-way ANOVA followed by Tukey's HSD post hoc test. $p < 0.05$ was considered statistically significant. (*) $p < 0.05$, (**) $p < 0.01$, (***) $p < 0.001$, (****) $p < 0.0001$.

Figure S4. HFD feeding induced changes in myeloid cells following CDI. Bar plots showing frequencies of (a) CD11b$^+$ myeloid cells (b) CD206$^-$CD11c$^+$ and CD206$^+$CD11c$^-$ macrophages in the colonic lamina propria from ND-fed and HFD-fed male and female mice following CDI at 3 and 6 dpi (n = 5-8). Radar plots showing scaled mean of frequencies of (c) myeloid cell subsets, and (d) dendritic cell subsets in antibiotic-treated (mock) and CDI male and female mice at 3 and 6 dpi (n = 5-8). Data in bar plots are presented as mean ± SEM. Statistical significance was assessed by two-way ANOVA followed by Tukey's HSD post hoc test. $p < 0.05$ was considered statistically significant. (*) $p < 0.05$, (**) $p < 0.01$, (***) $p < 0.001$, (****) $p < 0.0001$.

Figure S5. Changes in ILCs during acute and recovery phase following CDI. Radar plots showing scaled mean of frequencies of (a) ILC subsets, (b) cytokines produced by ILCs in antibiotic-treated (mock) and CDI male and female mice at 3 and 6 dpi (n = 5-8). (c) Bar plots showing frequencies of $CD127^{+}CD90^{+}$ innate lymphoid cell subsets (ILCs: T-bet$^{+}$ ILC1s, GATA-3$^{+}$ ILC2s and Rorγt$^{+}$ ILC3s); and (d) cytokine-production in ILCs (IFN-γ$^{+}$ ILC1s, IL13$^{+}$ ILC2s, and IL22$^{+}$ ILC3s) in the colonic lamina propria from ND- and HFD-fed male and female mice following CDI at 3 and 6 dpi (n=5-8). Data in bar plots are presented as mean ± SEM. Statistical significance was assessed by two-way ANOVA followed by Tukey's HSD post hoc test. $p < 0.05$ was considered statistically significant. (*) $p < 0.05$, (**) $p < 0.01$, (***) $p < 0.001$, (****) $p < 0.0001$.

Figure S6. HFD fed mice exhibited impaired microbiota restoration following CDI. Bar plots showing mean relative abundance of bacterial taxa identified in cecal samples from ND- and HFD-fed males and females at 6 dpi following CDI. Pink asterisks indicate statistically significant differences between ND-fed and HFD-fed female mice, and black asterisks indicate statistically significant differences between ND-fed and HFD-fed male mice. Statistical significance was determined by Wilcoxon test. $p < 0.05$ was considered statistically significant. (*) $p < 0.05$, (**) $p < 0.01$, (***) $p < 0.001$, (****) $p < 0.0001$.

Figure S7. HFD fed mice showed temporal changes in gene expression following CDI. (a) Heatmap showing k-means clustering ($k = 3$) and scaled expression of the differentially expressed genes (DEGs) in cecal samples, identified from the three-way interaction between diet, treatment, and dpi among ND-fed and HFD-fed male and female antibiotic-treated and CDI mice at 3 dpi and 6 dpi. (b) Gene ontology (GO) analysis performed using Enrichr showing the top 10 enriched terms for each cluster; node size and color represent FDR values.

Figure S8. HFD-fed male mice showed distinct gene expression changes following CDI. Volcano plots showing significantly upregulated and downregulated genes between ND-fed and HFD-fed male mice at (a) 3 dpi and (b) 6 dpi. (c-h) Expression patterns of unique and shared DEGs identified by Venn analysis using genes from ND-fed and HFD-fed male mice at (a) 3 dpi and (b) 6 dpi.

Figure S9. Heatmap showing scaled expression of genes involved in inflammation-, interferon-, cell death-, T cell mediated-mechanisms in ND- and HFD-fed male and female antibiotic-treated and CDI mice at 3 and 6 dpi.

Figure S10. Heatmap showing correlations between module eigengenes and bacterial taxa identified by weighted gene co-expression network analysis (WGCNA) in ND-fed and HFD-fed male and female in mock and CDI mice at 3 and 6 dpi.

Figure S1

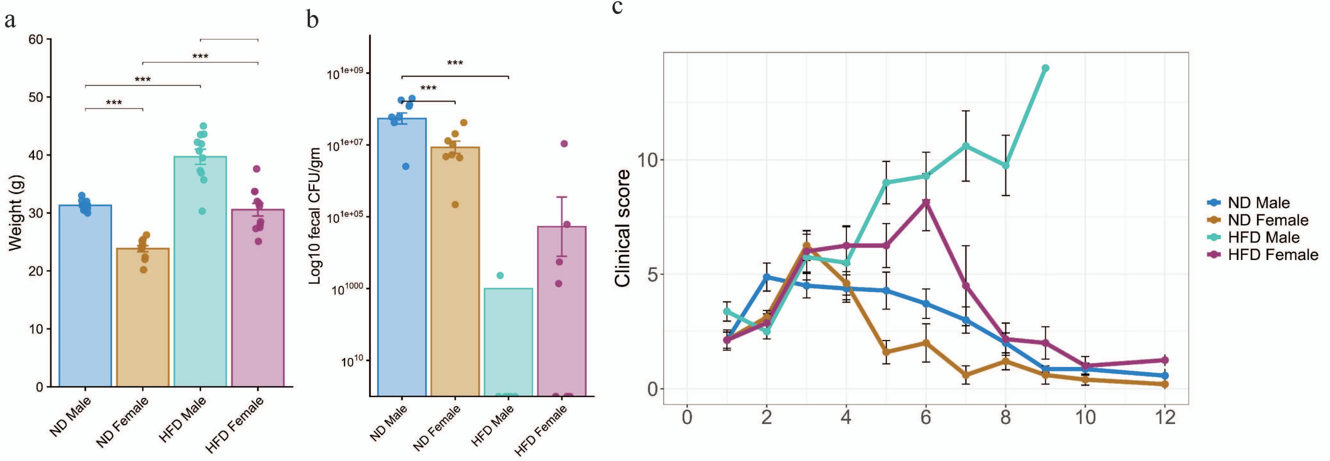

a
b
c
Weight (g)
***
ND Male
ND Female
HFD Male
HFD Female
Log10 fecal CFU/gm
Clinical score
ND Male
ND Female
HFD Male
HFD Female

Figure S2

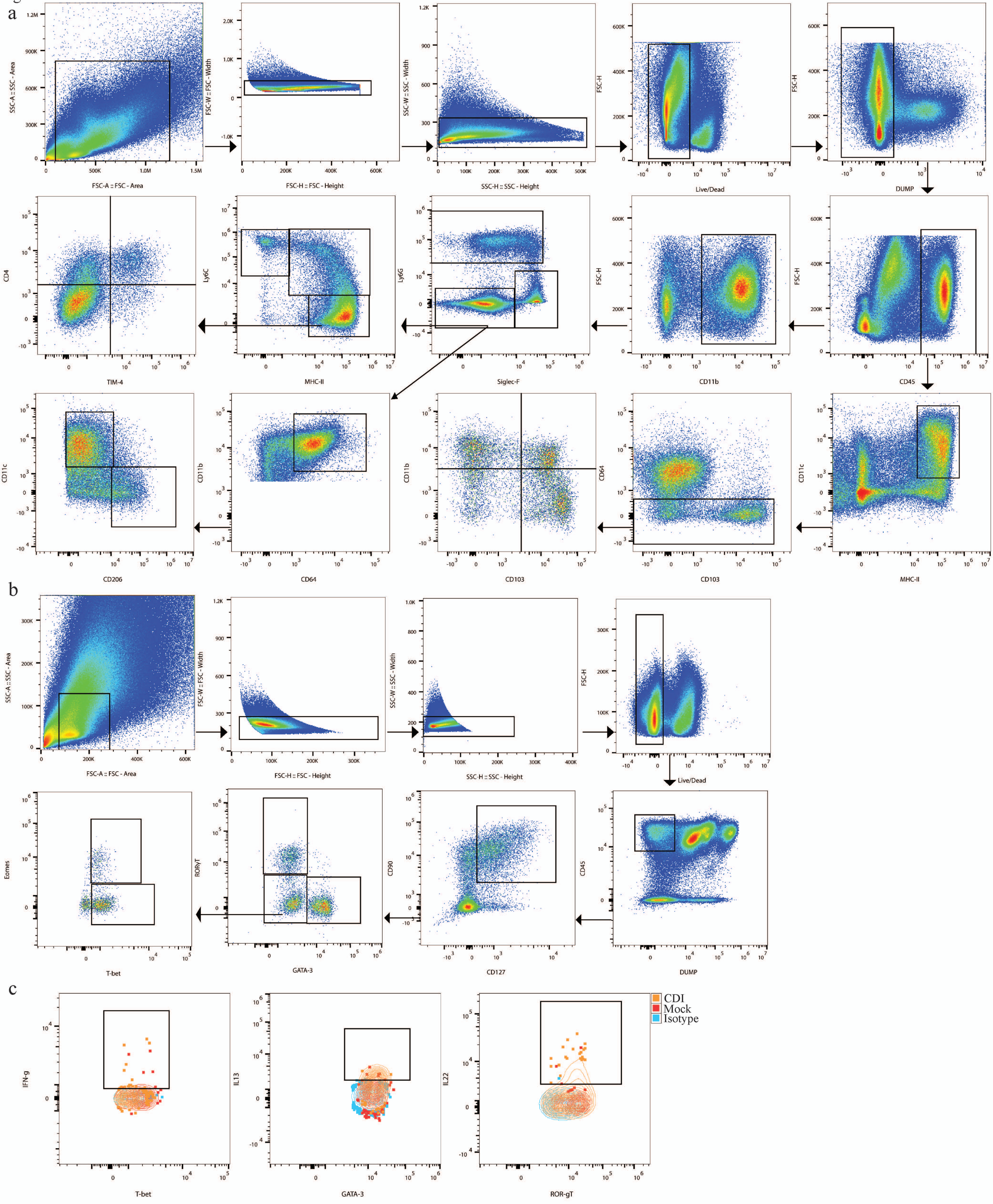
a
SSC-A :: SSC - Area
FSC-A :: FSC - Area
FSC-W :: FSC - Width
FSC-H :: FSC - Height
SSC-W :: SSC - Width
SSC-H :: SSC - Height
FSC-H
Live/Dead
DUMP
CD4
TIM-4
Ly6C
MHC-II
Ly6G
Siglec-F
CD11b
CD45
CD11c
CD206
CD64
CD103
b
SSC-A :: SSC - Area
FSC-A :: FSC - Area
FSC-W :: FSC - Width
FSC-H :: FSC - Height
SSC-W :: SSC - Width
SSC-H :: SSC - Height
Live/Dead
Eomes
T-bet
RORγT
GATA-3
CD90
CD127
CD45
DUMP
c
CDI
Mock
Isotype
IFN-g
T-bet
IL13
GATA-3
IL22
ROR-gT

Figure S3

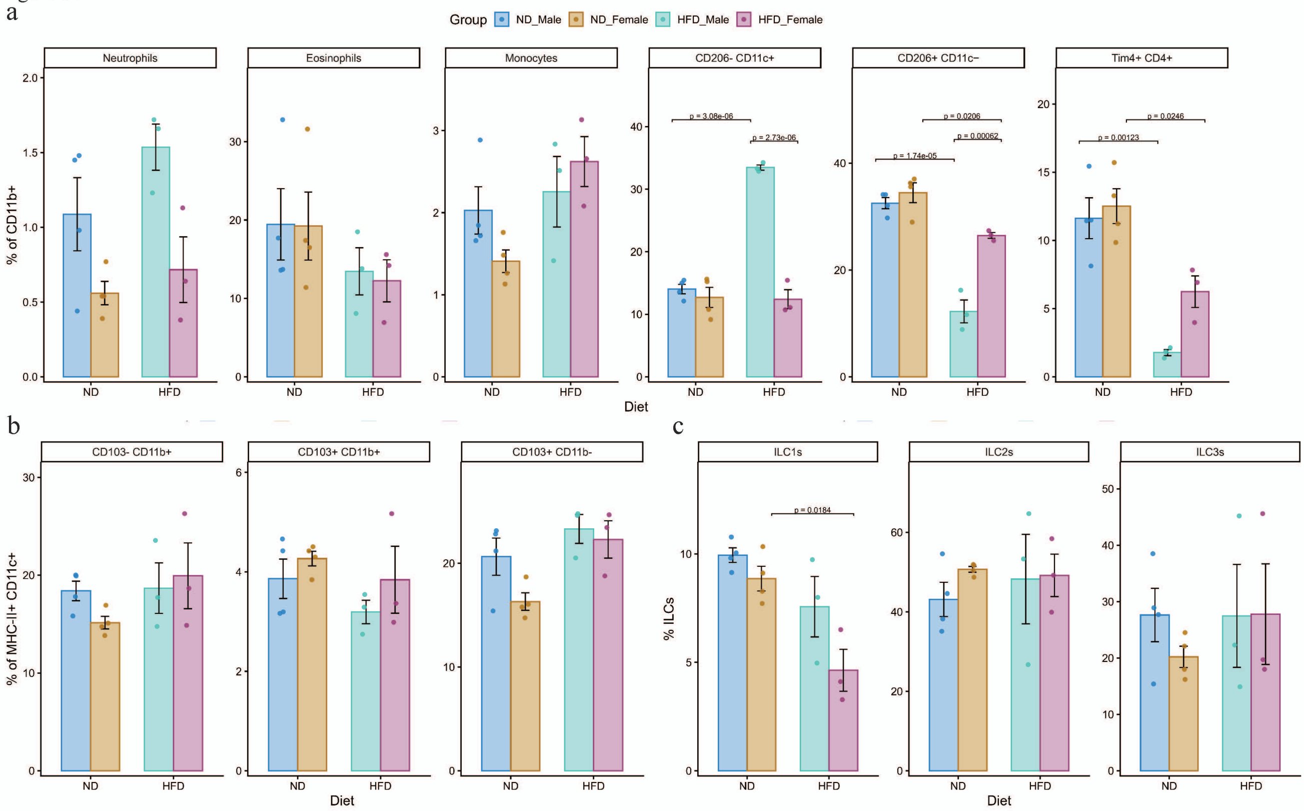
a
Group ND_Male ND_Female HFD_Male HFD_Female
Neutrophils
Eosinophils
Monocytes
CD206- CD11c+
CD206+ CD11c−
Tim4+ CD4+
% of CD11b+
p = 3.08e-06
p = 2.73e-06
p = 0.0206
p = 0.00062
p = 1.74e-05
p = 0.0246
p = 0.00123
ND
HFD
Diet
b
CD103- CD11b+
CD103+ CD11b+
CD103+ CD11b-
% of MHC-II+ CD11c+
c
ILC1s
ILC2s
ILC3s
% ILCs
p = 0.0184

Figure S4

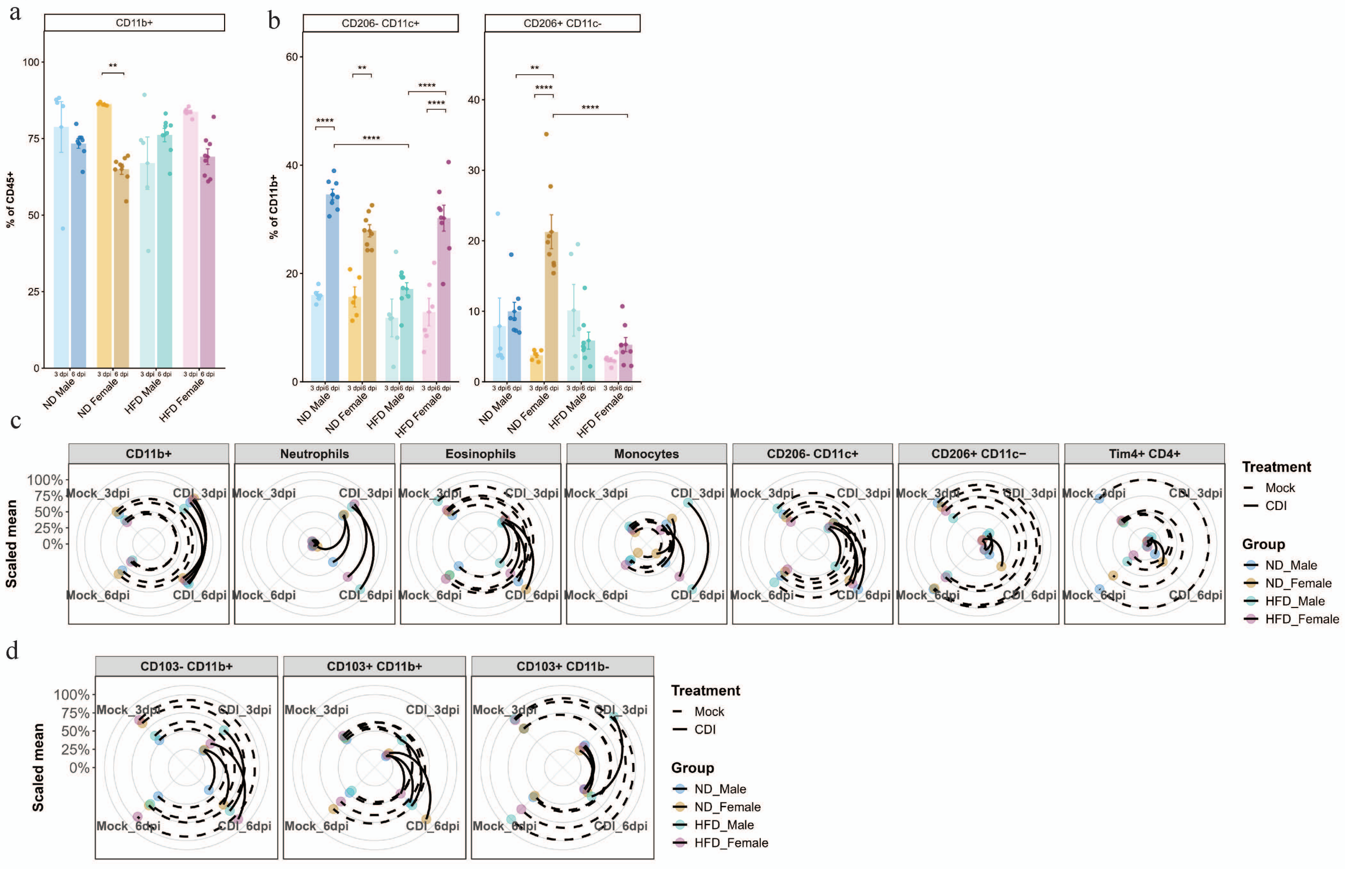
a
CD11b+
% of CD45+
ND Male
ND Female
HFD Male
HFD Female
3 dpi
6 dpi
b
CD206- CD11c+
CD206+ CD11c-
% of CD11b+
c
CD11b+
Neutrophils
Eosinophils
Monocytes
CD206- CD11c+
CD206+ CD11c−
Tim4+ CD4+
Scaled mean
Mock_3dpi
CDI_3dpi
Mock_6dpi
CDI_6dpi
Treatment
Mock
CDI
Group
ND_Male
ND_Female
HFD_Male
HFD_Female
d
CD103- CD11b+
CD103+ CD11b+
CD103+ CD11b-

Figure S5

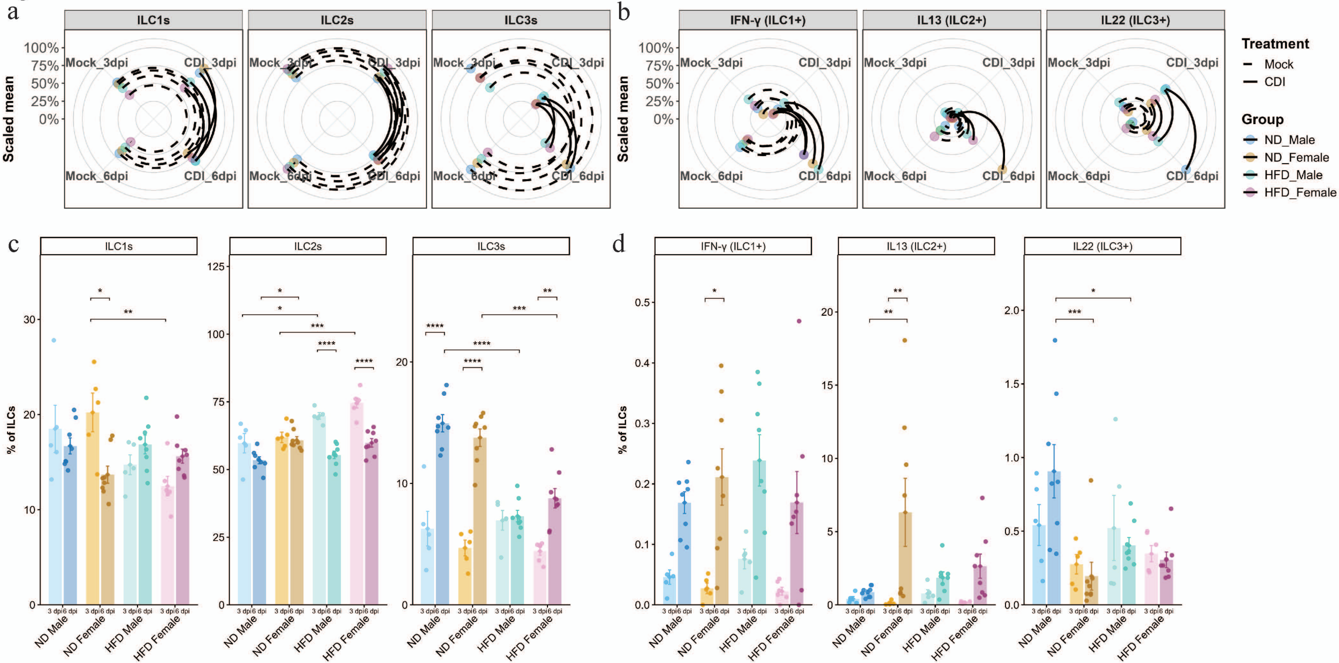
a
ILC1s
ILC2s
ILC3s
Scaled mean
100%
75%
50%
25%
0%
Mock_3dpi
CDI_3dpi
Mock_6dpi
CDI_6dpi
b
IFN-γ (ILC1+)
IL13 (ILC2+)
IL22 (ILC3+)
Treatment
Mock
CDI
Group
ND_Male
ND_Female
HFD_Male
HFD_Female
c
d
% of ILCs
3 dpi 6 dpi
ND Male
ND Female
HFD Male
HFD Female

Figure S6

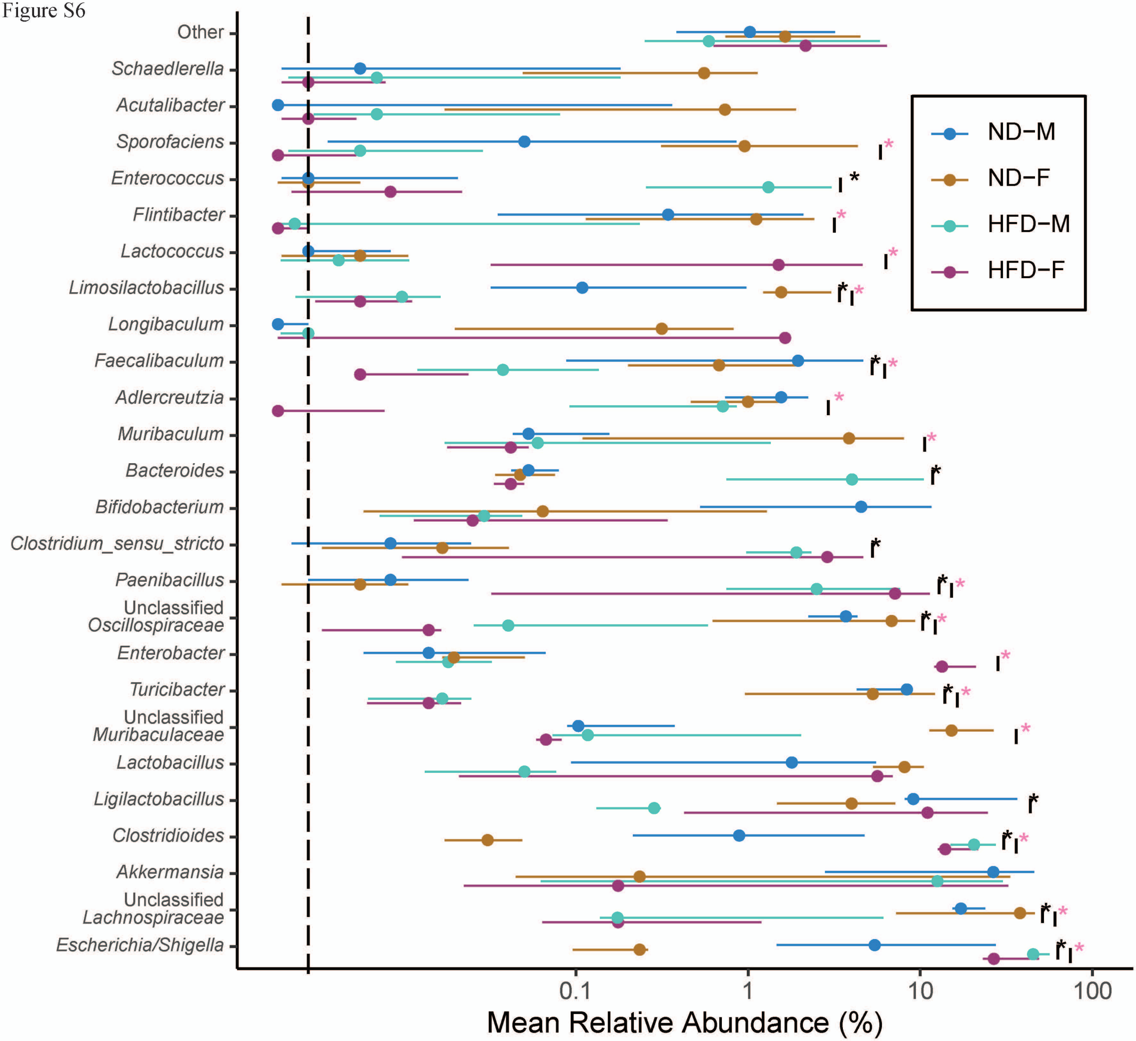

Other
Schaedlerella
Acutalibacter
Sporofaciens
Enterococcus
Flintibacter
Lactococcus
Limosilactobacillus
Longibaculum
Faecalibaculum
Adlercreutzia
Muribaculum
Bacteroides
Bifidobacterium
Clostridium_sensu_stricto
Paenibacillus
Unclassified Oscillospiraceae
Enterobacter
Turicibacter
Unclassified Muribaculaceae
Lactobacillus
Ligilactobacillus
Clostridioides
Akkermansia
Unclassified Lachnospiraceae
Escherichia/Shigella
ND−M
ND−F
HFD−M
HFD−F
0.1
1
10
100
Mean Relative Abundance (%)

Figure S7

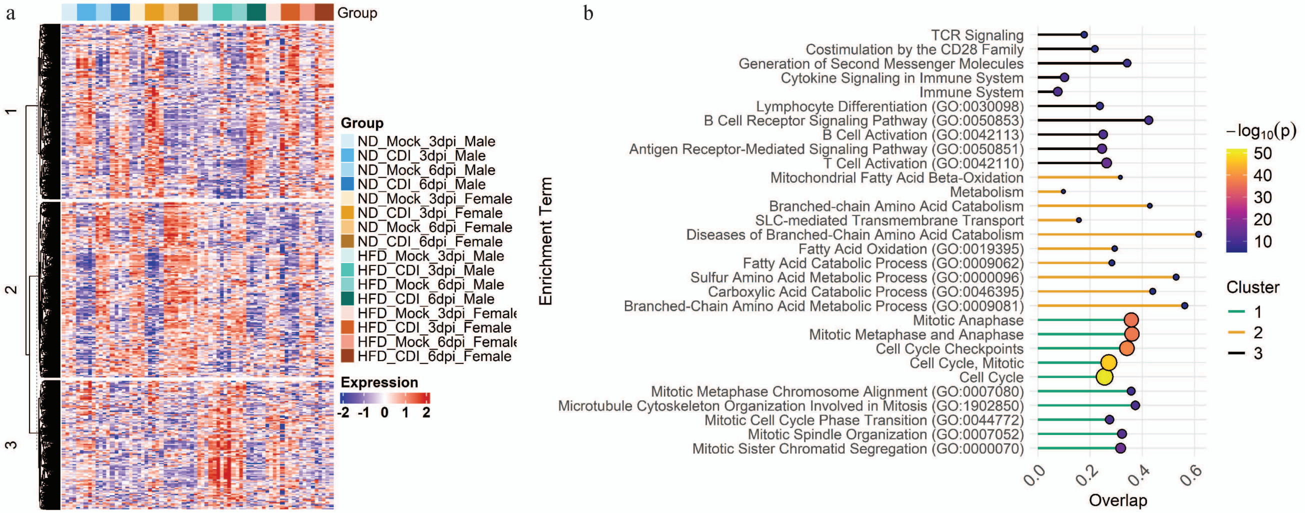
a
Group
1
2
3
Group
ND_Mock_3dpi_Male
ND_CDI_3dpi_Male
ND_Mock_6dpi_Male
ND_CDI_6dpi_Male
ND_Mock_3dpi_Female
ND_CDI_3dpi_Female
ND_Mock_6dpi_Female
ND_CDI_6dpi_Female
HFD_Mock_3dpi_Male
HFD_CDI_3dpi_Male
HFD_Mock_6dpi_Male
HFD_CDI_6dpi_Male
HFD_Mock_3dpi_Female
HFD_CDI_3dpi_Female
HFD_Mock_6dpi_Female
HFD_CDI_6dpi_Female
Expression
-2 -1 0 1 2
b
Enrichment Term
TCR Signaling
Costimulation by the CD28 Family
Generation of Second Messenger Molecules
Cytokine Signaling in Immune System
Immune System
Lymphocyte Differentiation (GO:0030098)
B Cell Receptor Signaling Pathway (GO:0050853)
B Cell Activation (GO:0042113)
Antigen Receptor-Mediated Signaling Pathway (GO:0050851)
T Cell Activation (GO:0042110)
Mitochondrial Fatty Acid Beta-Oxidation
Metabolism
Branched-chain Amino Acid Catabolism
SLC-mediated Transmembrane Transport
Diseases of Branched-Chain Amino Acid Catabolism
Fatty Acid Oxidation (GO:0019395)
Fatty Acid Catabolic Process (GO:0009062)
Sulfur Amino Acid Metabolic Process (GO:0000096)
Carboxylic Acid Catabolic Process (GO:0046395)
Branched-Chain Amino Acid Metabolic Process (GO:0009081)
Mitotic Anaphase
Mitotic Metaphase and Anaphase
Cell Cycle Checkpoints
Cell Cycle, Mitotic
Cell Cycle
Mitotic Metaphase Chromosome Alignment (GO:0007080)
Microtubule Cytoskeleton Organization Involved in Mitosis (GO:1902850)
Mitotic Cell Cycle Phase Transition (GO:0044772)
Mitotic Spindle Organization (GO:0007052)
Mitotic Sister Chromatid Segregation (GO:0000070)
0.0 0.2 0.4 0.6
Overlap
−log10(p)
50
40
30
20
10
Cluster
1
2
3

Figure S8

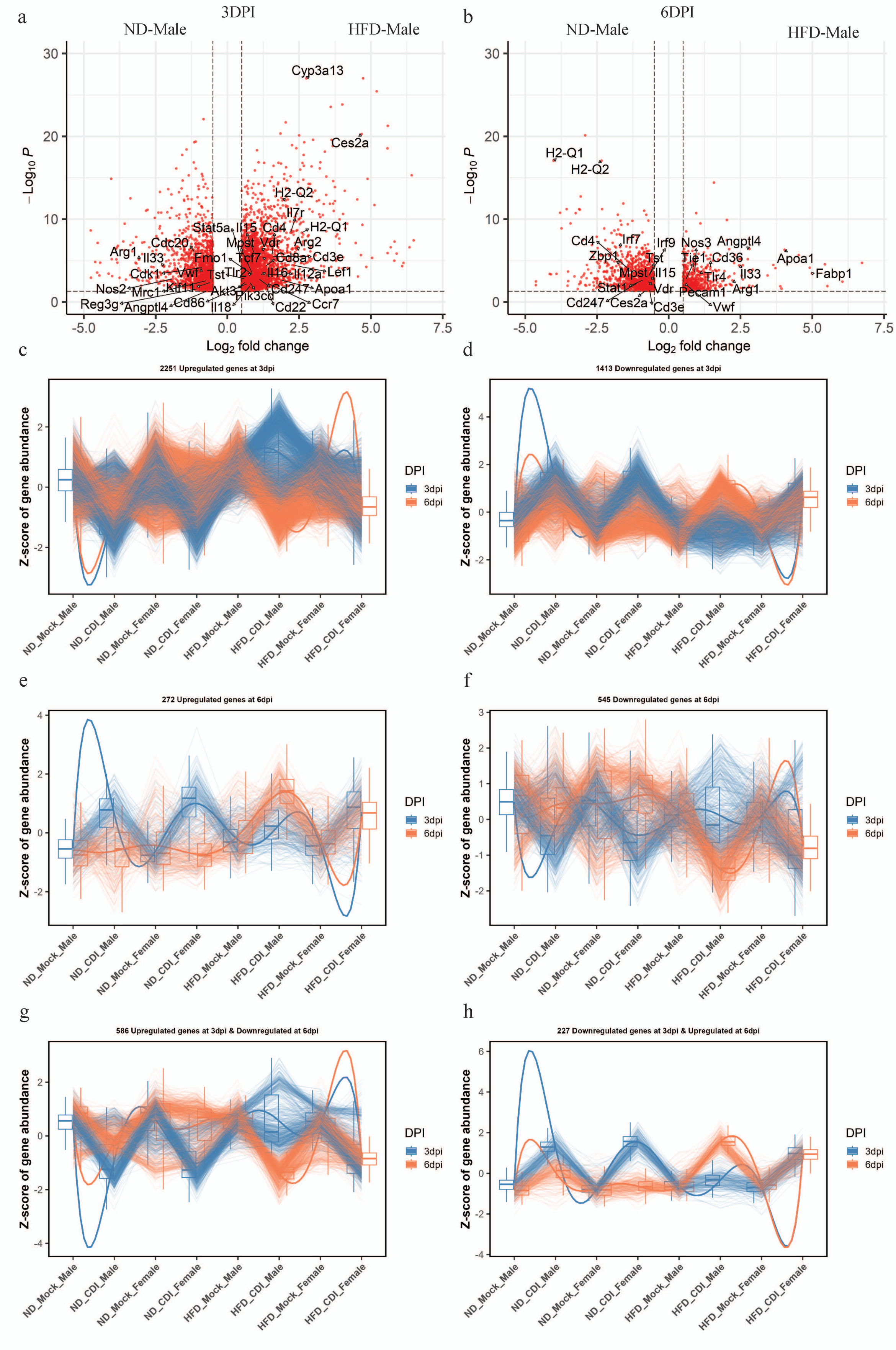
a
3DPI
ND-Male
HFD-Male
b
6DPI
ND-Male
HFD-Male
c
2251 Upregulated genes at 3dpi
d
1413 Downregulated genes at 3dpi
e
272 Upregulated genes at 6dpi
f
545 Downregulated genes at 6dpi
g
586 Upregulated genes at 3dpi & Downregulated at 6dpi
h
227 Downregulated genes at 3dpi & Upregulated at 6dpi
Z-score of gene abundance
DPI
3dpi
6dpi
ND_Mock_Male
ND_CDI_Male
ND_Mock_Female
ND_CDI_Female
HFD_Mock_Male
HFD_CDI_Male
HFD_Mock_Female
HFD_CDI_Female

Figure S9

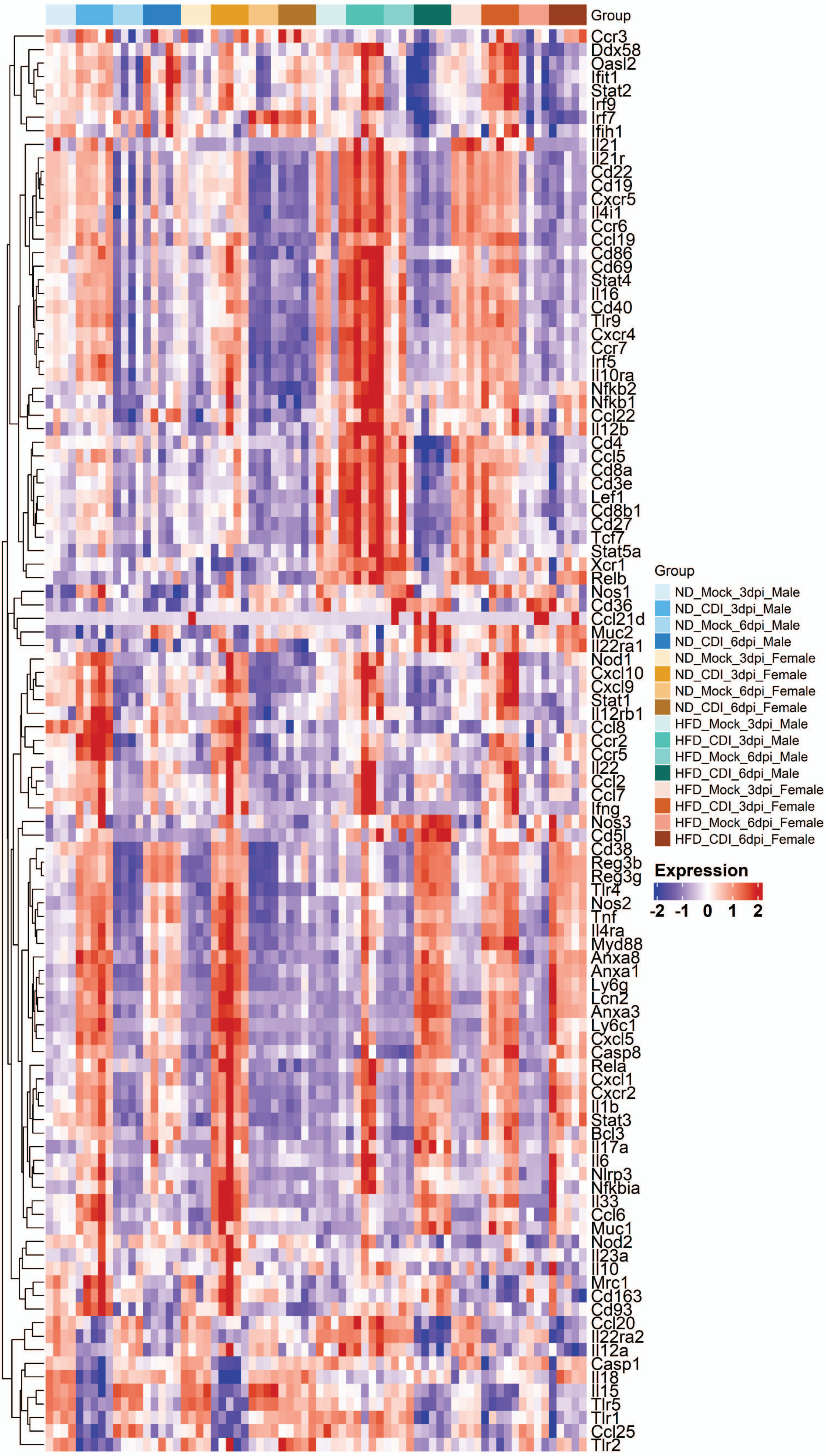
Group
Ccr3
Ddx58
Oasl2
Ifit1
Stat2
Irf9
Irf7
Ifih1
Il21
Il21r
Cd22
Cd19
Cxcr5
Il4i1
Ccr6
Ccl19
Cd86
Cd69
Stat4
Il16
Cd40
Tlr9
Cxcr4
Ccr7
Irf5
Il10ra
Nfkb2
Nfkb1
Ccl22
Il12b
Cd4
Ccl5
Cd8a
Cd3e
Lef1
Cd8b1
Cd27
Tcf7
Stat5a
Xcr1
Relb
Nos1
Cd36
Ccl21d
Muc2
Il22ra1
Nod1
Cxcl10
Cxcl9
Stat1
Il12rb1
Ccl8
Ccr2
Ccr5
Il22
Ccl2
Ccl7
Ifng
Nos3
Cd5l
Cd38
Reg3b
Reg3g
Tlr4
Nos2
Tnf
Il4ra
Myd88
Anxa8
Anxa1
Ly6g
Lcn2
Anxa3
Ly6c1
Cxcl5
Casp8
Rela
Cxcl1
Cxcr2
Il1b
Stat3
Bcl3
Il17a
Il6
Nlrp3
Nfkbia
Il33
Ccl6
Muc1
Nod2
Il23a
Il10
Mrc1
Cd163
Cd93
Ccl20
Il22ra2
Il12a
Casp1
Il18
Il15
Tlr5
Tlr1
Ccl25
Tlr2
Group
ND_Mock_3dpi_Male
ND_CDI_3dpi_Male
ND_Mock_6dpi_Male
ND_CDI_6dpi_Male
ND_Mock_3dpi_Female
ND_CDI_3dpi_Female
ND_Mock_6dpi_Female
ND_CDI_6dpi_Female
HFD_Mock_3dpi_Male
HFD_CDI_3dpi_Male
HFD_Mock_6dpi_Male
HFD_CDI_6dpi_Male
HFD_Mock_3dpi_Female
HFD_CDI_3dpi_Female
HFD_Mock_6dpi_Female
HFD_CDI_6dpi_Female
Expression
-2
-1
0
1
2

Figure S10

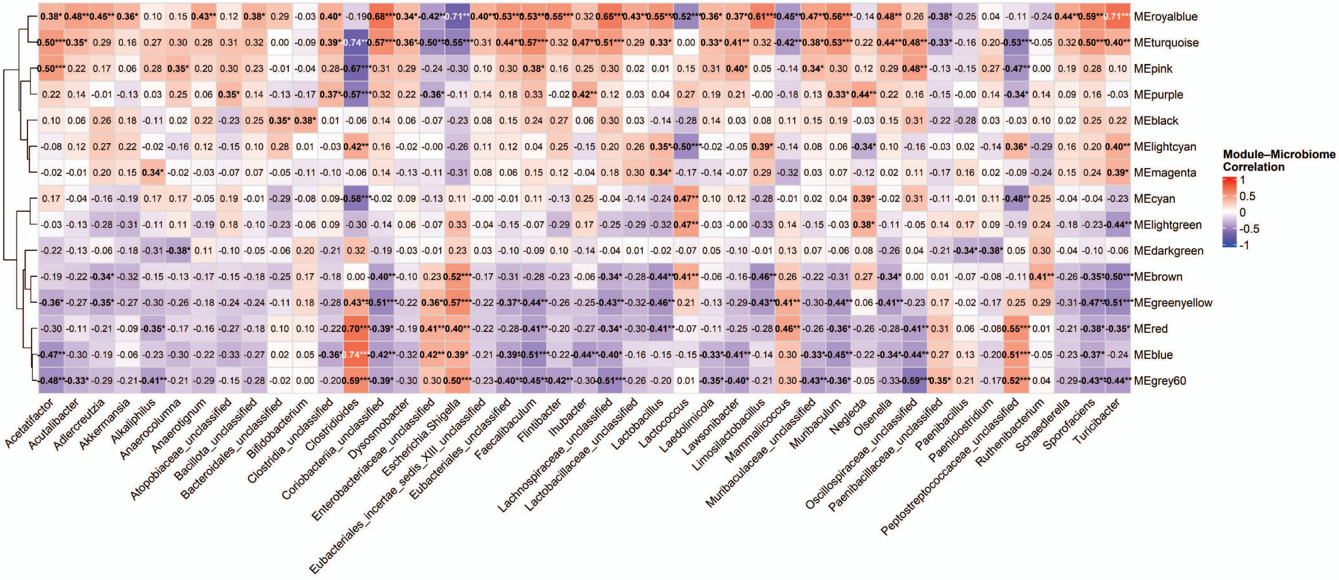
MEroyalblue
MEturquoise
MEpink
MEpurple
MEblack
MElightcyan
MEmagenta
MEcyan
MElightgreen
MEdarkgreen
MEbrown
MEgreenyellow
MEred
MEblue
MEgrey60
Module–Microbiome Correlation
1
0.5
0
-0.5
-1
Acetatifactor
Acutalibacter
Adlercreutzia
Akkermansia
Alkaliphilus
Anaerocolumna
Anaerotignum
Atopobiaceae_unclassified
Bacillota_unclassified
Bacteroidales_unclassified
Bifidobacterium
Clostridia_unclassified
Clostridioides
Coriobacteriia_unclassified
Dysosmobacter
Enterobacteriaceae_unclassified
Escherichia.Shigella
Eubacteriales_incertae_sedis_XIII_unclassified
Eubacteriales_unclassified
Faecalibaculum
Flintibacter
Ihubacter
Lachnospiraceae_unclassified
Lactobacillaceae_unclassified
Lactobacillus
Lactococcus
Laedolimicola
Lawsonibacter
Limosilactobacillus
Mammaliicoccus
Muribaculaceae_unclassified
Muribaculum
Neglecta
Olsenella
Oscillospiraceae_unclassified
Paenibacillaceae_unclassified
Paenibacillus
Paeniclostridium
Peptostreptococcaceae_unclassified
Ruthenibacterium
Schaedlerella
Sporofaciens
Turicibacter
